# Rhythmic temporal structure organizes recurrent dynamics to support sequential working memory

**DOI:** 10.64898/2026.05.20.726720

**Authors:** Yitao Qin, Yun Yang, Qizhi Yang, Qinglai Wei, Tielin Zhang

**Author notes:** **For correspondence:** (TZ). These authors contributed equally to this work (Yitao Qin, Yun Yang, Qizhi Yang).

## Abstract

Rhythmic temporal structure improves working memory, but how this benefit emerges from recurrent dynamics remains unclear. Here, we trained excitatory–inhibitory recurrent neural networks with short-term synaptic plasticity to perform a sequential delayed match-to-sample task with either regular or jittered sample timing. Rhythmic input produced a small but reliable improvement in task accuracy and was associated with more differentiated population trajectories during encoding. This behavioral advantage was accompanied by an organization of population dynamics around the dominant input frequency: temporal regularity progressively brought stimulus arrivals closer to preferred encoding phases, modulated phase advancement during stimulus presentation, and reduced the deviation of inter-stimulus phase-progression frequency from the dominant input rhythm. As a result, internal oscillations increasingly tracked the temporal structure of the input across the sequence, providing a phase-based scaffold for encoding ordered information. This scaffold preferentially supported temporal-order representations rather than uniformly enhancing all stimulus features. Decoding analyses further showed that stronger temporal regularity increased the fidelity and persistence of stimulus information in both neuronal activity and synaptic efficacy, whereas perturbing synaptic efficacy produced the largest impairment during delay-period maintenance. These findings suggest that rhythmic input supports sequential working memory by imposing a reliable temporal structure on recurrent dynamics and stabilizing synaptic-state representations.

## Introduction

Working memory is commonly characterized as a limited-capacity system for the temporary storage and manipulation of information required for ongoing cognition (***Baddeley, 2012***). In sequential working-memory tasks, this general requirement is extended to information distributed across successive events: individual items must be retained together with their ordinal or positional context. Research on memory for serial order has shown that order information is a major issue in verbal, visual, and spatial short-term memory, indicating that sequence memory cannot be fully explained by item storage alone (***Hurlstone et al., 2014***). Because sequence items are encountered successively, the temporal organization of the input stream may influence when item representations are encoded, how adjacent items are separated, and how ordered information is subsequently maintained.

Rhythm is defined as the serially ordered pattern of time intervals in a stimulus sequence (***Fiveash et al., 2022***). In our study, rhythmicity is operationalized as the temporal regularity of stimulus onsets, quantified by the variability of inter-onset intervals (IOIs). Under this definition, a sequence with identical or near-identical IOIs has stronger rhythmic structure, whereas greater IOI variability corresponds to weaker rhythmicity. Regular temporal structure can support temporal expectations by making the timing of upcoming events more predictable (***Large and Jones, 1999***; ***Nobre and Van Ede, 2018***). Consistent with this view, temporal expectation and rhythmic temporal structure have been shown to improve sensory processing, recognition memory, visual working-memory-guided behavior, and auditory working memory (***Rohenkohl et al., 2012***; ***Jones and Ward, 2019***; ***Jin et al., 2020***; ***Tian et al., 2025***). These findings establish that temporal regularity can benefit memory-related behavior, but they do not specify how such regularity is expressed in recurrent neural dynamics during sequential working memory.

One neural route through which temporal regularity may influence sequence encoding is the phase organization of low-frequency activity. Low-frequency oscillations provide a temporal scale over which neuronal excitability can fluctuate, whereas phase organization refers to the systematic relationship between external event timing and the phase state of ongoing neural activity (***Lakatos et al., 2005***; ***Schroeder and Lakatos, 2009***). In rhythmic sensory contexts, the phase of neural activity can become aligned with the temporal structure of external stimulation, thereby biasing processing toward expected moments of input (***Lakatos et al., 2008***; ***Obleser and Kayser, 2019***). Related work in working memory further shows that temporal expectations can dynamically prioritize visual working-memory representations, and that rhythmic temporal coordination of neural activity can reduce representational conflict among maintained items (***van Ede et al., 2017***; ***Abdalaziz et al., 2023***). These findings motivate the question of whether temporally regular input organizes recurrent population dynamics during sequence encoding, particularly by constraining the relationship between stimulus arrival and oscillatory phase state.

Sequential working memory also requires task-relevant information to persist across a stimulus-free delay. Two classes of neural mechanisms have been proposed to support such maintenance. Persistent-activity accounts emphasize sustained neuronal firing during delay periods, whereas activity-silent accounts propose that information can also be maintained in latent states that are not continuously expressed in elevated spiking activity (***Stokes, 2015***; ***Rose et al., 2016***; ***Barbosa et al., 2020***). Short-term synaptic plasticity provides a candidate synaptic mechanism for latent maintenance, because transient changes in synaptic efficacy can preserve information over short timescales without requiring continuous persistent firing (***Mongillo et al., 2008***; ***Masse et al., 2019***; ***Kozachkov et al., 2022***). Thus, it is necessary to examine both encoding-period population dynamics and delay-period maintenance in neuronal activity and synaptic efficacy for understanding how temporal regularity facilitates sequential working memory.

To explore the possible neural mechanism of how temporal regularity facilitates sequential working memory and eventually improves task performance, we trained a biologically constrained excitatory–inhibitory recurrent neural network with short-term synaptic plasticity to perform a sequential delayed match-to-sample task (***Song et al., 2016***; ***Barak, 2017***; ***Masse et al., 2019***; ***Kozachkov et al., 2022***). Following the task design of a previous study (***Tian et al., 2025***), our network model encoded a sequence of six oriented stimuli, maintained the sequence across a delay, and then judged whether a subsequent test sequence matched the sample sequence. We varied the temporal regularity of the sample sequence by comparing inputs with fixed versus variable IOIs while preserving the same sequence-memory requirement. This framework allowed us to examine how input temporal regularity shapes recurrent neural dynamics during different task periods.

## Results

### Rhythmic temporal structure improves task performance across task regimes

To mechanistically simulate the computational properties of biological neural systems, we constructed a biologically constrained recurrent neural network (RNN) (***Song et al., 2016***; ***Masse et al., 2019***). The network consists of an input layer with 30 direction-tuned neurons and 1 rule-tuned neuron, projecting to a recurrent layer comprising 80 excitatory and 20 inhibitory neurons (see Methods: Network models), adhering to the excitatory-inhibitory balance found in the mammalian cortex (***Wang, 2002***) (***Figure 1***A). A key computational feature of the network is the dynamic modulation of recurrent connection weights, implemented via short-term synaptic plasticity (STSP) (***Tsodyks and Markram, 1997***; ***Mongillo et al., 2008***) (see Methods: Short-term synaptic plasticity).

**Figure 1.**
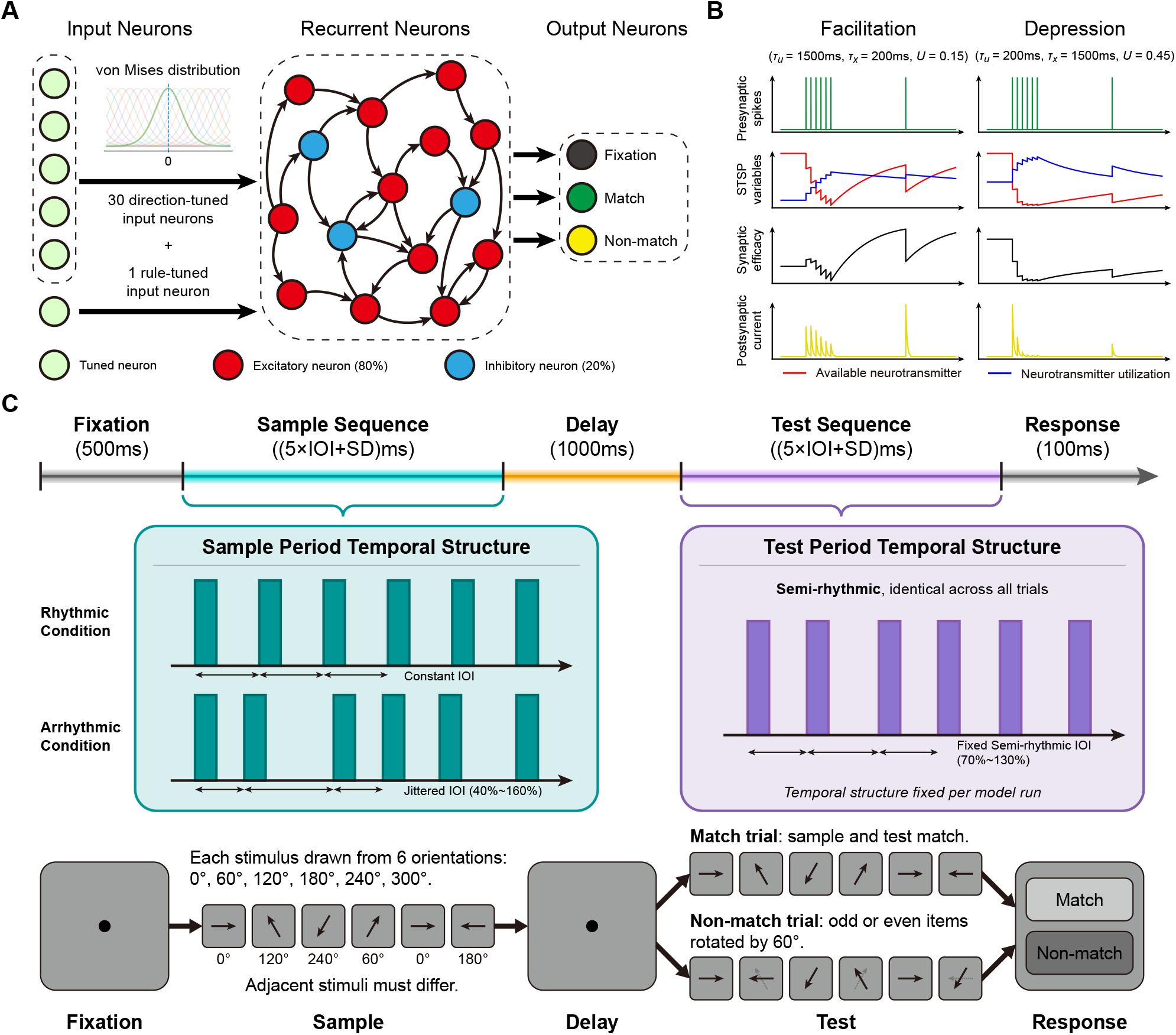
Network architecture, short-term synaptic plasticity (STSP) mechanism and sequence working memory task paradigm. (A) Schematic of the recurrent neural network (RNN) architecture. Sensory inputs from 30 tuned neurons are projected into a recurrent layer consisting of 80 excitatory (red) and 20 inhibitory (blue) neurons. The recurrent layer then projects to output units representing different decision states (fixation, match, non-match). (B) Microscopic dynamics of STSP. The evolution of the available neurotransmitter fraction (*x*, red line) and the utilization fraction (*u*, blue line) in response to presynaptic spikes (green) is illustrated for both short-term facilitation (left) and short-term depression (right). The transient synaptic efficacy (*u* ⋅ *x*, black line) dynamically determines the amplitude of the resulting postsynaptic current (yellow). (C) Timeline of the sequence working memory task. Following a 500 ms fixation period, a sequence of six oriented bar stimuli is presented during the sample period, followed by a 1000 ms delay. In the rhythmic condition, sample items are presented with constant inter-onset intervals (equal IOIs). In the arrhythmic condition, the sequence features temporal jitter with variable IOIs. (when stimulus duration (SD) = 100 ms, IOI ∈ {250, 300, 350, 400, 450, 500} ms; when SD = 120 ms, IOI ∈ {300, 350, 400, 450, 500} ms; when SD = 140 ms, IOI ∈ {350, 400, 450, 500} ms) The test sequence is semi-rhythmic in both conditions, prompting a final match/non-match behavioral response.

The synaptic efficacy of neuron *j* at time *t* is defined as the product of the fraction of available neurotransmitter, *x*_*j*_(*t*), and the utilization fraction, *u*_*j*_(*t*): *S*_*j*_(*t*) = *x*_*j*_(*t*) ⋅ *u*_*j*_(*t*). Consequently, the total synaptic input received by neuron *i* is 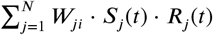, where *W*_*ji*_ denotes the baseline connection weight from neuron *j* to neuron *i*, and *R*_*j*_(*t*) represents the instantaneous firing rate of neuron *j*.

Previous research indicates that temporal expectation and structure not only modulate perceptual gain but also significantly influence working memory performance and memory-guided action execution (***van Ede et al., 2017***; ***Jin et al., 2020***). To examine the robustness of this rhythmic advantage within our neural network model, we designed a Delayed Match-to-Sample (DMS) task (***Ullén et al., 2014***) (***Figure 1***C). In this task, the network is required to determine, during the final response period, whether a stimulus sequence presented in the test period is identical to that in the sample period (***Miller et al., 1996***).

We manipulated the temporal structure of the sample period using two primary conditions:

1. Rhythmic condition: Six stimuli were presented consecutively with a constant Inter-Onset Interval (IOI).
2. Arrhythmic condition: Temporal jitter was introduced to the constant IOI, while ensuring no temporal overlap between stimuli to maintain physical discriminability (***Grahn and Brett, 2007***; ***Snyder et al., 2024***; ***Levitin et al., 2018***).

To ensure that performance differences arose from sample-period encoding rather than test-period timing, the test sequence was presented with a semi-rhythmic temporal structure that was identical across conditions for each model instance. For sample construction, a “match” trial was defined as exact directional correspondence between sample and test sequences; a “non-match” trial involved a 60^◦^ clockwise rotation of stimuli at either odd or even positions (***Tian et al., 2025***). As an additional control, we also tested a position-unconstrained variant in which the changed stimuli could occur at arbitrary sequence positions. Comparable performance in this setting argues against a decision strategy based solely on the earliest stimulus items.

To quantify the impact of rhythmic input structure on working memory, we evaluated the performance of 100 independently trained networks across varying temporal scales. While the networks performed proficiently in both conditions, statistical analysis revealed a small but reliable improvement in sequence-level accuracy under rhythmic input compared with arrhythmic input. As shown in ***Figure 2***A, rhythmic input produced a small but reliable accuracy improvement in match trials (rhythmic: 0.991 ± 0.002; arrhythmic: 0.987 ± 0.002; mean ± s.e.m.; two-sided paired t-test, *n* = 100 independent networks, *p* < 10^−3^). A similar advantage was observed in “non-match” trials involving a 60^◦^ rotation (rhythmic: 0.973 ± 0.006 vs. arrhythmic: 0.969 ± 0.007, mean ± s.e.m., *p* < 10^−3^, *n* = 100 independent networks trained with SD in {100, 120, 140} ms, two-tailed paired t-test). These results are consistent with the idea that temporal regularity can support sequence-level performance in this working-memory task.

**Figure 2.**
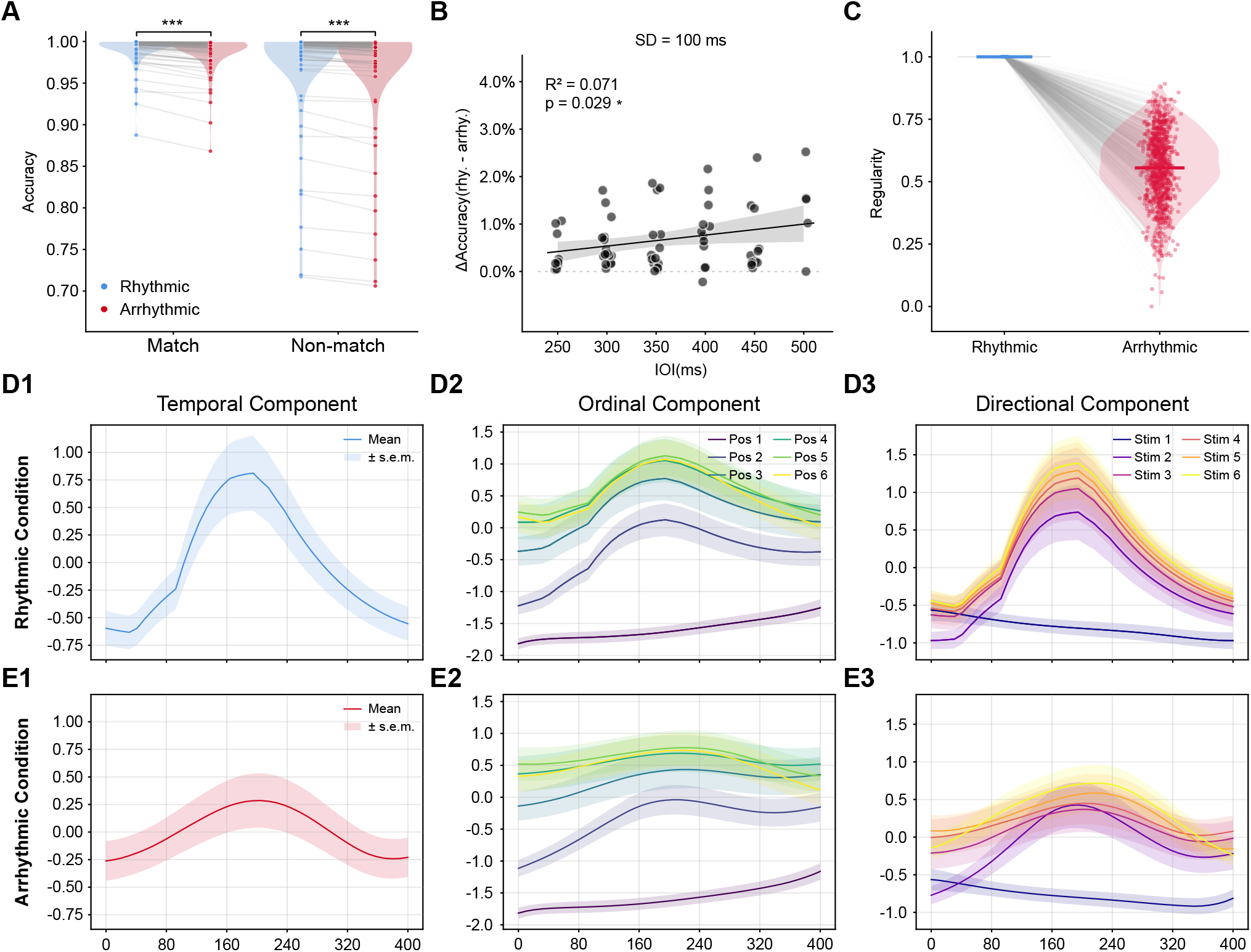
Rhythmic temporal structure improves sequence memory and organizes population-level representations. (A) Behavioral accuracy in the working memory task under rhythmic and arrhythmic conditions (match and non-match trials). Rhythmic sequences showed significantly higher task accuracy (two-sided paired t-test, *n* = 100 independent networks, *p* < 0.001). (B) The behavioral benefit of rhythmicity was positively correlated with the base IOI (Pearson correlation, two-sided, *n* = 67 independent networks with *SD* = 100 ms, *p* < 0.05). (C) Distribution of the trial regularity index (*R*) under rhythmic and arrhythmic conditions. Larger R values indicate a higher degree of rhythmicity (see Methods: Task paradigm). (D-E) Population-level demixed principal component analysis (dPCA) of network representations under rhythmic (D1-D3) and arrhythmic (E1-E3) conditions. Rhythmic input were associated with more separable low-dimensional population trajectories across temporal, ordinal, and stimulus-related components. Unless otherwise stated, data are shown as mean ± s.e.m. Statistical tests are two-sided where applicable. Significance levels are denoted as \**p* < 0.05, \*\**p* < 0.01, and \*\*\**p* < 0.001.

Furthermore, we tested whether the accuracy difference between rhythmic and arrhythmic conditions depended on IOI length. We fixed the stimulus duration (SD = 100 ms) while varying the mean sequence IOI, and calculated the accuracy difference between conditions (Δ*Accuracy* = *Acc*_*Rhy*_ − *Acc*_*Arr*_). Regression analysis revealed a significant positive correlation between the rhythmic accuracy gain and IOI length (Pearson correlation, *R*^2^ = 0.071, *p* = 0.029, *n* = 67 independent networks trained with SD = 100 ms; ***Figure 2***B). The positive correlation remained consistent across networks trained with SD = 120 and 140 ms. The rhythmic advantage tended to be larger at longer IOIs, suggesting that temporal regularity may become more beneficial when longer temporal gaps between successive items place greater demands on working-memory maintenance.

### Rhythmic input reorganizes task representations in low-dimensional geometry

To reveal how rhythmic input supports superior working memory at the neural population level, we investigated the geometric structure of the network’s internal representations. Given that population neural activity is typically conceptualized as low-dimensional dynamical processes, and dimensionality reduction methods can effectively characterize and compare these dynamical structures (***Cunningham and Yu, 2014***; ***Vyas et al., 2020***), we employed demixed principal component analysis (dPCA) to reveal the model’s representational structure (***Kobak et al., 2016***). We cropped and aligned the neural activity for each stimulus during the sample period across 10 independent networks (IOI = 400 ms) and projected it into low-dimensional subspaces. This approach enabled isolation and visualization of the network’s dynamical trajectories within three distinct component subspaces: temporal component, ordinal component, and directional component.

Across all three task-relevant subspaces, we calculated the proportion of variance *R*_*m*_ explained by the PC axes (see Methods: Demixed principal component analysis). The first demixed principal component (dPC1) consistently captured the vast majority of the signal variance (time subspace: 65.45%; position subspace: 55.40%; stimulus subspace: 58.20%), while the explained variance of subsequent components decreased substantially (dPC2 was consistently below 24%). This steep decline indicates that a dominant component captured a large fraction of task-related variance within each subspace.

We first examined the temporal component, which revealed how the network internally tracks the passage of time (***Buonomano and Maass, 2009***; ***Paton and Buonomano, 2018***). Under rhythmic conditions, the network exhibited robust, large-amplitude temporal trajectories that peaked during stimulus presentation (***Figure 2***D1). These trajectories suggest stronger temporal modulation of the population state under rhythmic input. In contrast, trajectory amplitudes appeared attenuated under arrhythmic conditions (***Figure 2***E1).

Subsequent projection into the ordinal subspace showed more visually separated trajectories across sequence position, consistent with reduced overlap among adjacent item representations (***Figure 2***D2). These visualizations suggest greater overlap among adjacent sequence-position trajectories under arrhythmic input (***Figure 2***E2).

Lastly, in the directional component subspace, which captures specific directional features, rhythmic input expands the separation between stimulus-specific population trajectories, despite identical physical input intensity across conditions (***Figure 2***D3). This population-level geometric expansion is consistent with improved population-level discriminability (***Chung and Abbott, 2021***), whereas such trajectory separation was markedly reduced under arrhythmic conditions (***Figure 2***E3).

Together, these results suggest that rhythmic input is associated with stronger low-dimensional separability of temporal, ordinal, and stimulus-related population trajectories.

### How the temporal structure of external input organizes neural oscillatory patterns in recurrent neural networks

Temporal structure is an important dimension in how cognitive systems process external input. Previous behavioral and neuroscience studies have shown that temporal regularity in external events can support temporal attention and temporal expectation, allowing processing resources to be preferentially allocated at expected moments and improving sensory processing and behavioral performance (***Large and Jones, 1999***; ***Nobre and Van Ede, 2018***). At the neural mechanism level, temporal expectation is thought to be closely related to the phase alignment of low-frequency neural oscillations to external rhythms. This alignment can periodically modulate the excitability of local neural populations, allowing high-excitability phases to align more stably with the occurrence of expected events, thereby enhancing responses to relevant inputs (***Lakatos et al., 2005, 2008***; ***Arnal and Giraud, 2012***). We therefore examined whether regular temporal input organized oscillatory dynamics in the trained recurrent networks. To address this question, it is first necessary to operationally define the strength of temporal regularity in the input. Building on the previous binary task conditions (rhythmic vs. arrhythmic), we further re-quantified rhythmic strength as a continuous variable *R* (***Figure 2***C), defined as *R* = (*R*_*raw*_ − *min*(*R*_*raw*_) / max(*R*_*raw*_) − min(*R*_*raw*_)), 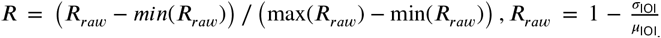 (see Methods: Trial-wise temporal regularity and stimulation frequency). When *R* = 1, the external input is presented uniformly in time; if the timing of stimulus occurrence within a single trial is more irregular, then *R* is smaller.

External rhythms can entrain low-frequency neural oscillations, enabling neural activity to form phase organization and selective responses on timescales matched to the stimulus rhythm (***Lakatos et al., 2008***; ***Schroeder and Lakatos, 2009***). Therefore, we first examined whether population activity in the trained RNN showed frequency-resolved modulation around the dominant input frequency *f*_0_ (see Methods: Trial-wise temporal regularity and stimulation frequency), which characterizes the average rate of the external temporal sequence. To test whether the network’s activity responses exhibited frequency selectivity, we calculated the response strength of network neuronal activity across different frequency bands (see Methods: Spectral analysis of population activity). The results showed that significant stimulus-responsive activity was not uniformly distributed across frequencies, but was most prominent near the dominant input frequency *f*_0_ (***Figure 3***A1). Moreover, nearly all analyzed frequency bands were sensitive to the temporal structure of the external input (***Figure 3***A2), indicating that the temporal structure of the external input systematically influenced the network’s oscillatory patterns. Further spectral analysis showed that neural responses within the *f*_0_ band increased with increasing regularity *R* (***Figure 3***B), indicating that the effect of temporal regularity first manifested as the selective organization of oscillatory components at a specific timescale. Based on this result, we defined the neighborhood of *f*_0_ as the target band for subsequent analyses.

**Figure 3.**
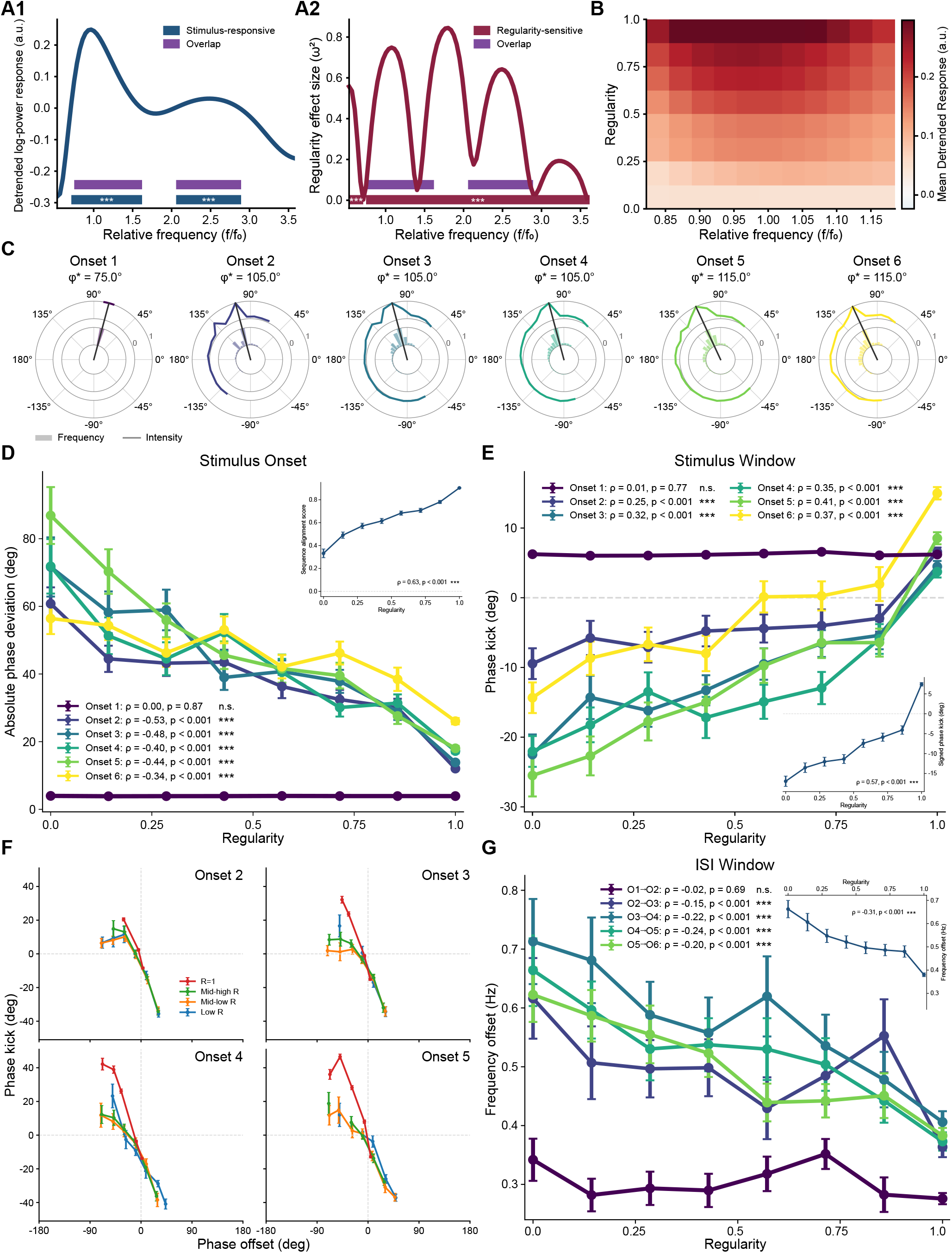
Temporal regularity selectively organizes sample-period oscillatory dynamics. (A1) Frequency-resolved response strength *T* (*f*) of population activity during the sample epoch, plotted as a function of relative frequency (*f* /*f*_0_). The solid line denotes the across-trial mean response (*n* = 4000 trials). Blue horizontal bars indicate frequency clusters showing significant stimulus-evoked responses, assessed using a one-sample sign-flip cluster-permutation test against zero (*t* statistic, *B* = 1000 permutations). Significant stimulus-responsive clusters were observed at 0.71–1.58 *f* /*f*_0_ (1.78–3.96 Hz; Cohen’s *d*_*z*_ = 1.74, cluster-level *p* < 0.001) and 2.05–2.85 *f* /*f*_0_ (5.14–7.14 Hz; Cohen’s *d*_*z*_ = 0.97, cluster-level *p* < 0.001). Purple bars indicate the portions of stimulus-responsive frequencies that overlapped with significant regularity-sensitive clusters. (A2) Regularity-dependent modulation of spectral responses across frequency. The curve denotes the frequency-wise effect size *ω*^2^ from a one-way ANOVA across regularity bins. Red horizontal bars indicate significant regularity-sensitive clusters, assessed using label-permutation cluster testing on *F* statistics (*B* = 1000 permutations). Significant regularity-sensitive clusters were observed at 0.53–0.68 *f* /*f*_0_ (1.33–1.69 Hz; peak/mean *ω*^2^ = 0.53/0.38, cluster-level *p* < 0.001) and 0.75–3.50 *f* /*f*_0_ (1.87–8.75 Hz; peak/mean *ω*^2^ = 0.80/0.41, cluster-level *p* < 0.001). Purple bars indicate overlap between stimulus-responsive and regularity-sensitive frequency ranges. (B) Regularity-dependent response strength around the retained stimulation-frequency band. Each row corresponds to one regularity bin, and color indicates the bin-averaged response strength *T* (*f*) in arbitrary units. Directional modulation was quantified by Spearman’s rank correlation between regularity-bin medians and bin-averaged responses (*ρ* = 0.65, permutation test, *p* < 0.001), indicating stronger *f*_0_-band responses with increasing temporal regularity. (C) Preferred encoding phase for each sample stimulus position. Polar plots show the distribution of stimulus-arrival phases and their associated local response profiles. Grey bars denote phase-frequency histograms, colored curves denote density-regularized local response scores, and black vectors indicate the preferred phase 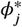, defined as the phase-bin center with the maximal density-regularized local response score for each stimulus position. (D) Temporal regularity reduces the deviation between stimulus-arrival phase and preferred encoding phase. The absolute circular phase deviation *D*_*j*_ between the stimulus-arrival phase and 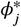 is plotted as a function of temporal regularity for each stimulus position. The inset shows the across-position summary of phase alignment as a function of regularity. Statistical annotations report Spearman’s rank correlations between trial-wise temporal regularity and phase deviation. (E) Temporal regularity modulates stimulus-window phase kick. Phase kick *K*_*j*_ was defined as the circular difference between the observed phase advance during the stimulus window and the nominal phase advance expected from *f*_0_. Positive values indicate faster-than-*f*_0_ phase advancement during stimulus presentation, whereas negative values indicate slower-than-*f*_0_ advancement. The inset shows the across-position summary of stimulus-window phase kick as a function of regularity. Statistical annotations report Spearman’s rank correlations between trial-wise temporal regularity and phase kick. (F) Phase kick as a function of stimulus-arrival phase offset. Phase offset was defined as the signed circular distance between the stimulus-arrival phase and the preferred encoding phase 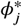. Negative offsets indicate that the stimulus arrived before the preferred phase, whereas positive offsets indicate that it arrived after the preferred phase. Colors denote temporal-regularity groups (*R* = 1, mid-high *R*, mid-low *R*, and low *R*). Dashed lines indicate zero phase offset and zero phase kick. (G) Temporal regularity reduces the deviation of ISI phase progression from the dominant input frequency. The plotted metric is the absolute frequency deviation | *IF*_ISI_ − *f*_0_ |, which quantifies how far the ongoing phase-advance rate deviated from the dominant temporal scale of the input. Smaller values indicate that the internal phase advanced at a rate closer to *f*_0_. The first transition was not reliably modulated by regularity (O1→O2, *ρ* = −0.02, *p* = 0.69), whereas later transitions showed significant negative correlations with regularity (O2→O3, *ρ* = −0.15; O3→O4, *ρ* = −0.22; O4→O5, *ρ* = −0.24; O5→O6, *ρ* = −0.20; all *p* < 0.001). The inset shows the sequence-level summary across ISI windows (*ρ* = −0.31, *p* < 0.001). Unless otherwise stated, points indicate regularity-bin means and error bars denote mean ± s.e.m. Statistical tests are two-sided where applicable. Significance levels are denoted as \**p* < 0.05, \*\**p* < 0.01, and \*\*\**p* < 0.001. For permutation-based cluster tests, significance refers to cluster-level *p* values.

When the cognitive task begins, the network enters a preparatory state during the fixation period, and the neural activity at this time reflects the network’s spontaneous oscillatory dynamics; the presentation of external stimulus input perturbs this oscillatory pattern. Previous studies have shown that regular temporal input can periodically reorganize the excitability state of neural populations by entraining the phase of low-frequency neural oscillations, thereby allowing high-excitability phases to align more stably with the occurrence time of expected events (***Lakatos et al., 2008***; ***Schroeder and Lakatos, 2009***). Therefore, we next examined from the perspective of phase how stimulus arrival and the temporal interval pattern between stimuli affect oscillations within the network. Preferred phase refers to a specific phase position in the oscillatory cycle at which the system is in the highest excitability or lowest inhibition state. When an external stimulus falls at the preferred phase, the network response is strongest (***Lakatos et al., 2005***; ***Schroeder and Lakatos, 2009***). On this basis, we counted the phase distribution and oscillatory amplitude at the time of stimulus presentation for each sequence position across 4000 trials, and used phase-bin center with the maximal density-regularized response score as the preferred phase for that sequence position (***Figure 3***C). We then examined the absolute circular deviation between the actual phase at which each stimulus fell and its preferred phase under different levels of temporal regularity (see Methods: Oscillatory phase organization during sample encoding). The results showed that as the regularity of the external temporal input increased, the absolute deviation between the phase at which the stimulus fell and the preferred phase decreased significantly (***Figure 3***D). However, this negative correlation did not appear immediately at the beginning of the sequence (Onset 1, *ρ* = −0.00, *p* = 8.74 × 10^−1^), but became significant with the gradual accumulation of temporal context (Onset 2–6, all *p* < 0.001). This indicates that at the early stage of sequence presentation, the system had not yet accumulated sufficient temporal information to establish a stable internal temporal reference; with the continuous input of external temporal structure, temporal regularity gradually organized the phase structure of network oscillations.

The continuous presentation of stimuli imposes sustained perturbations on oscillations within the network. At this point, does the change in oscillatory phase structure simply reflect passive resetting driven by the stimulus, or is it also accompanied by regulation of the current phase state by the network? Under ideal conditions without stimulus input and without considering internal network noise, oscillations in the *f*_0_ band should advance uniformly along their corresponding ideal periodic phase structure, and the effect of stimulus input is reflected in the deviation of the actual phase trajectory. Based on this, we turned off the internal noise of the network, quantified stimulus-window phase advancement through fitting an amplitude-weighted robust linear slope to the unwrapped phase trajectory within each stimulus window. Phase kick was defined as the circular difference between the observed phase advance and the nominal phase advance expected from the retained *f*_0_ band (see Methods: Oscillatory phase organization during sample encoding). Positive values therefore indicate faster-than-*f*_0_ phase advancement during stimulus presentation, whereas negative values indicate slower-than-*f*_0_ advancement. The results showed that stimulus presentation caused *f*_0_ oscillations to exhibit accelerated advancement already at the beginning of the sequence (***Figure 3***E, Onset 1), and with increasing temporal regularity of the external input, this acceleration effect in phase advancement was further enhanced for Onset 2–6 (all *p* < 0.001), whereas weak temporal regularity of the external input caused the advancing speed of oscillatory phase organization to lag behind its original speed. If the phase at the onset of stimulus presentation lagged behind the preferred phase, then the average phase advancement speed over the subsequent stimulus presentation window would be greater than the original *f*_0_ advancement speed; conversely, the phase advancement speed would be relatively lower (***Figure 3***F). This indicates that the phase change during stimulus presentation is not entirely a passive response; the network is also actively adjusting the advancing speed of phase organization according to changes in the temporal structure of the external input, which may be helpful for promoting active correction of internal oscillations toward a state favorable for encoding.

During the interval from the offset of the current stimulus to the onset of the next stimulus, we next examined how the internal oscillatory phase continued to advance after stimulus-driven perturbation. To quantify phase progression during this inter-stimulus interval (ISI), we estimated the instantaneous frequency of the internal oscillation and computed its absolute deviation from the dominant input frequency, | *IF*_ISI_ − *f*_0_ | (see Methods: Inter-stimulus phase-progression frequency). This measure captures the magnitude of the difference between the ongoing phase-advance rate and the phase-advance rate expected at the dominant temporal scale of the input. At the earliest transition, this frequency deviation was not reliably modulated by temporal regularity (***Figure 3***G, O1→O2, *ρ* = −0.02, *p* = 0.69). However, as the sequence unfolded, the deviation between ISI phase-progression frequency and *f*_0_ decreased with increasing temporal regularity across later intervals (***Figure 3***G, O2→O3, *ρ* = −0.15, *p* < 0.001; O3→O4, *ρ* = −0.22, *p* < 0.001; O4→O5, *ρ* = −0.24, *p* < 0.001; O5→O6, *ρ* = −0.20, *p* < 0.001). The sequence-level summary showed the same negative relationship (*ρ* = −0.31, *p* < 0.001). Thus, after temporal context had accumulated, stronger input regularity reduced the deviation of ISI phase progression from the dominant input timescale, indicating that internal oscillations advanced more closely to *f*_0_ between successive stimuli.

Then, within the single event window from the presentation of the current stimulus to the start of the next stimulus, did a stable correspondence form between the internal phase and the external temporal structure? To address this, we constructed the external input phase according to the temporal interval pattern of the stimuli. For each stimulus interval, we defined an external phase that advanced linearly within the window according to the actual duration of that interval, such that it completed one full cycle between two adjacent stimuli. The purpose of doing this was to map stimulus intervals of different lengths onto the same phase coordinate system, thereby characterizing the progression state of the current external temporal structure so that it could be directly compared with the internal oscillatory phase of the network (see Methods: Coupling between external temporal phase and neural oscillatory phase). ***Figure 4***A shows the differences between the external input phase and the internal *f*_0_ oscillatory phase in two representative trials under different levels of temporal regularity. On this basis, we calculated the external phase and internal amplitude-weighted phase-locking value (PLV) across all IOI windows under different levels of temporal regularity. The results showed that the phase-locking strength within all IOIs increased with increasing temporal regularity *R* (***Figure 4***B, all *p* < 0.001), and this positive effect became stronger as the stimulus sequence unfolded. This indicates that the tracking of external input temporal structure by the network’s neural oscillations during the sample period is a process that is gradually formed and continuously maintained. Taken together, we can infer that the phase organization of internal neural oscillations by the temporal regularity of external input may be cumulatively formed by the following processes: during stimulus presentation, the network drives internal oscillations toward a phase state that is more favorable for encoding; during the inter-stimulus interval, the average frequency of internal oscillations gradually tunes toward the dominant frequency of the external input. As a result, within each IOI window, the phase-locking relationship between internal oscillations and external input events becomes stronger with increasing temporal regularity. Thus, the temporal regularity of external input does not influence network activity only at discrete stimulus moments, but continuously shapes the oscillatory dynamics of the network throughout the entire stimulus sequence.

**Figure 4.**
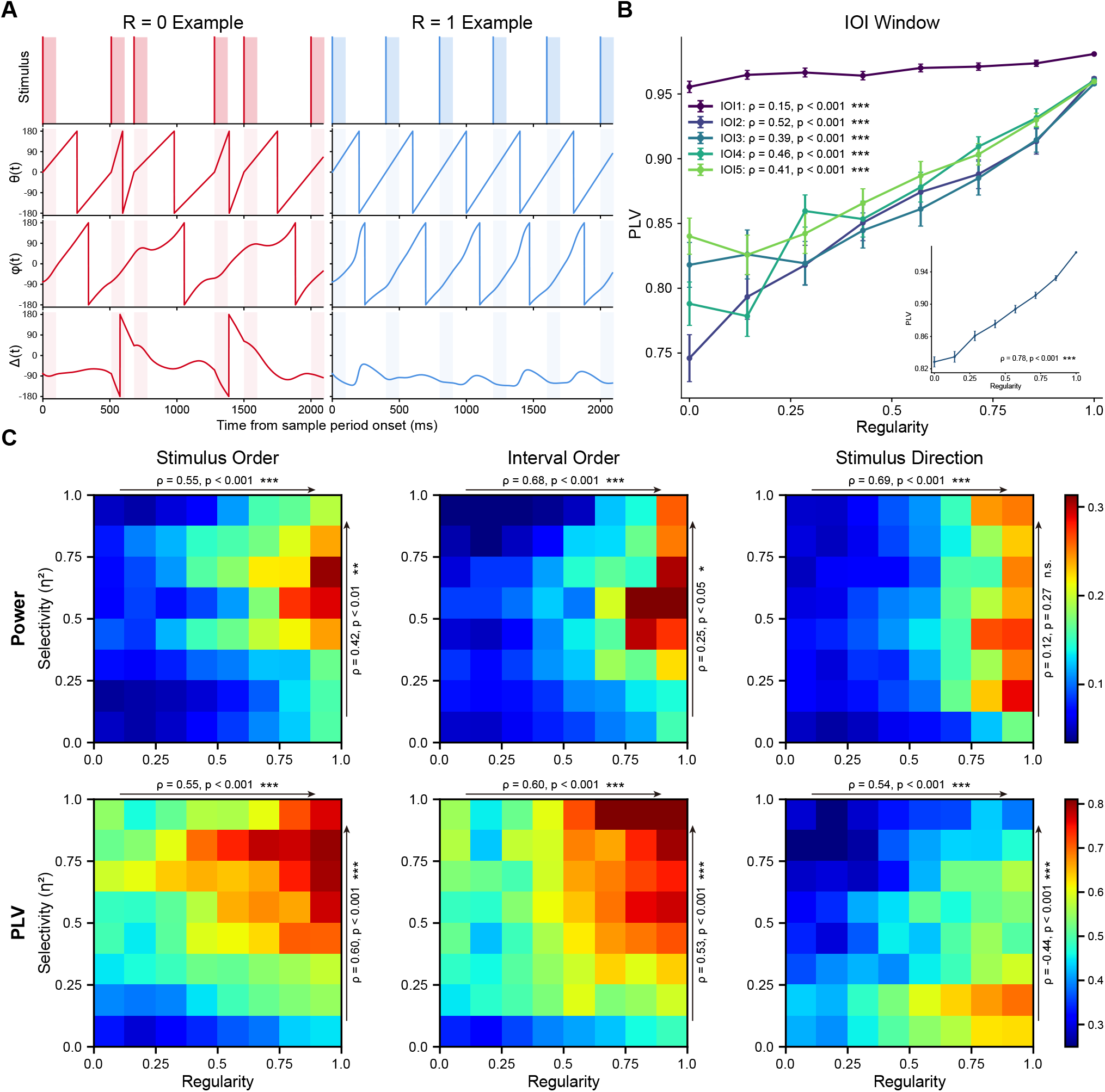
Oscillatory organization selectively supports temporal encoding during the sample period. (A) Representative trials illustrating the construction of external temporal phase and its relationship to internal oscillatory phase under low-regularity (*R* = 0) and perfectly regular (*R* = 1) input. Shaded regions indicate stimulus presentation windows. The external temporal phase *θ*(*t*) advances linearly from 0 to 2*π* within each inter-onset interval and resets at the next stimulus onset. The internal phase was obtained from the retained *f*_0_-band population signal using the same causal complex filter as in the phase-progression analyses. (B) Amplitude-weighted phase-locking value between internal neural phase and external temporal phase as a function of trial-wise temporal regularity. Points indicate regularity-bin means, and error bars denote mean ± s.e.m. across trials (*n* = 4000 trials). Statistical annotations report Spearman’s rank correlation between *R* and PLV_*j*_ for each IOI window. The inset shows the average PLV across IOI windows, 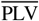, as a function of regularity, with the corresponding Spearman’s rank correlation (permutation test, *ρ* = 0.78, *P* < 0.001). (C) Joint heatmaps show the joint distribution of trial regularity (x-axis) and neuronal selectivity (*η*^2^, y-axis) for oscillatory power (top row) and phase-locking value (PLV; bottom row). Columns correspond to selectivity for stimulus order, interval order, and stimulus direction. The statistics shown above and to the right of each panel summarize the monotonic trends along the regularity and selectivity dimensions, respectively. Statistics indicate the median model-level Spearman’s rank correlation; significance was evaluated using Fisher’s *z* transformation followed by a sign-flip test. Unless otherwise stated, data are shown as mean ± s.e.m. Statistical tests are two-sided where applicable. Significance levels are denoted as \**p* < 0.05, \*\**p* < 0.01, and \*\*\**p* < 0.001.

### Neural oscillations preferentially support temporal-order encoding during the sample period

Neural oscillations are often considered a mechanism for organizing information in time, but their representational consequences are not necessarily uniform across different task variables. In sequential working memory, this distinction is particularly important: the same sample stream contains both temporal-ordinal information, such as when each item occurs and how intervals are ordered, and feature-specific information, such as stimulus direction. The preceding analyses showed that temporal regularity progressively organized the network’s endogenous rhythm, producing stronger stimulus-matched power and more stable phase alignment during the sample period. A remaining question is therefore whether this internally organized rhythm supports all forms of sample-period representation equally, or whether its functional contribution is biased toward intrinsically temporal variables.

To address this question, we next related trial-level regularity and oscillatory measures to the neuronal selectivity of different task variables using a matrix-based analysis (***Figure 4***C; see Methods: Selectivity analysis of oscillatory organization). Across all three variables, regularity showed robust positive associations with both power and PLV, indicating that increasing temporal regularity globally strengthened the internally matched oscillatory state of the network. Thus, external rhythmicity broadly enhanced both the amplitude and the phase consistency of the sample-period rhythm. However, this enhancement was not translated uniformly into stronger selectivity. Instead, the matrix structure revealed that the consequence of rhythmic organization depended on which aspect of the sample sequence was being encoded.

For stimulus order and interval order, both power and PLV tended to increase together with selectivity. This pattern suggests that rhythmic organization was preferentially linked to the encoding of temporal structure. Notably, the relationship was especially clear for PLV, suggesting that precise phase coordination, rather than oscillatory magnitude alone, is closely related to the emergence of order-selective responses. In other words, rhythmic input appear to do more than simply elevate the overall excitability of the network. By organizing when neurons respond relative to the internal oscillatory cycle, they create a temporal scaffold that is particularly favorable for representing the sequential structure of the sample stream (***Nobre and Van Ede, 2018***; ***Abdalaziz et al., 2023***; ***Duecker et al., 2024***). By contrast, stimulus direction exhibited a different profile. Although regularity still enhanced the oscillatory state itself, directional selectivity showed no reliable association with power and a significant negative association with PLV. This dissociation suggests that stronger rhythmic alignment does not uniformly sharpen all forms of sample-period coding. Rather, the internally organized rhythm appears to preferentially stabilize those aspects of the input that are inherently temporal, such as when items occur and how they are ordered, instead of directly amplifying feature-specific selectivity for every stimulus attribute. This result suggests that the representational consequences of rhythmic organization differed across task variables, consistent with recent evidence that phase-related firing need not directly preserve physical stimulus order (***Liebe et al., 2025***).

This interpretation also helps integrate the present matrix result with the preceding analyses. Earlier population-level dPCA analysis (***Figure 2***D and ***Figure 2***E) showed that rhythmic input increases the separability of task-related trajectories, whereas the phase analyses from ***Figure 3***, ***Figure 4***A, and ***Figure 4***B demonstrated that temporal regularity progressively aligns internal oscillations with external timing and stabilizes within-IOI locking. ***Figure 4***C refines this picture by showing that the most direct encoding consequence of such oscillatory organization falls on temporal-order variables. Importantly, this does not contradict the earlier improvement in directional population geometry. Instead, it could suggest a functional distinction in scale: at the level of oscillatory coordination and single-unit selectivity, rhythmicity most directly supports temporal structure, whereas at the population level the same temporal scaffold may still improve directional readout indirectly, possibly by reducing overlap among successive items and imposing a more orderly dynamical regime during encoding (***Stokes, 2015***; ***Murray et al., 2017***).

These results are difficult to explain by a purely global-gain account. While external regularity strengthens the network’s internal oscillatory state in general, the resulting functional benefit appears to be expressed primarily through temporal structuring, suggesting that internally organized oscillations may preferentially support the stabilization and differentiation of temporal-order structure during the sample period.

### Improved decodability in neural activity and synaptic efficacy indicates stronger memory-related representations

The preceding results indicate that rhythmic input induces precise frequency and phase locking during the sample period. We next asked whether rhythmicity-related phase organization was associated with improved readout of stimulus direction.

Based on the “activity-silent” framework of working memory, short-term memory content can be maintained implicitly in synaptic efficacy while being explicitly reflected in residual neuronal activity (***Mongillo et al., 2008***; ***Stokes, 2015***; ***Masse et al., 2019***). To test stimulus information decodability, we trained a linear Support Vector Machine (SVM) to decode the direction of each sequence stimulus from the population states of both neuronal activity and synaptic efficacy across time (***Meyers et al., 2008***; ***King and Dehaene, 2014***) (see Methods: Population decoding). To directly link phase organization to information readout, we performed a PLV-binned decoding analysis during the sample period, in which item-level trials were grouped into fixed PLV intervals and decoding accuracy was averaged within each bin (***Figure 5***A). Higher PLV was associated with higher decoding accuracy for both neuronal activity and synaptic efficacy, indicating that stronger phase locking tracked item-level decoding accuracy. To further establish the relationship between information decodability and external temporal statistical structure, we then performed a parametric regression analysis linking the mean decoding accuracy with the trial-level regularity index *R* (***Figure 5***B1 and ***Figure 5***B2). The results indicate a significant positive correlation between *R* and the decoding accuracy of both neuronal activity and synaptic efficacy across sample and delay periods. This suggests that temporal regularity is associated with stronger decodable stimulus information in both neuronal activity and synaptic efficacy, consistent with a dual-state interpretation of working-memory maintenance.

**Figure 5.**
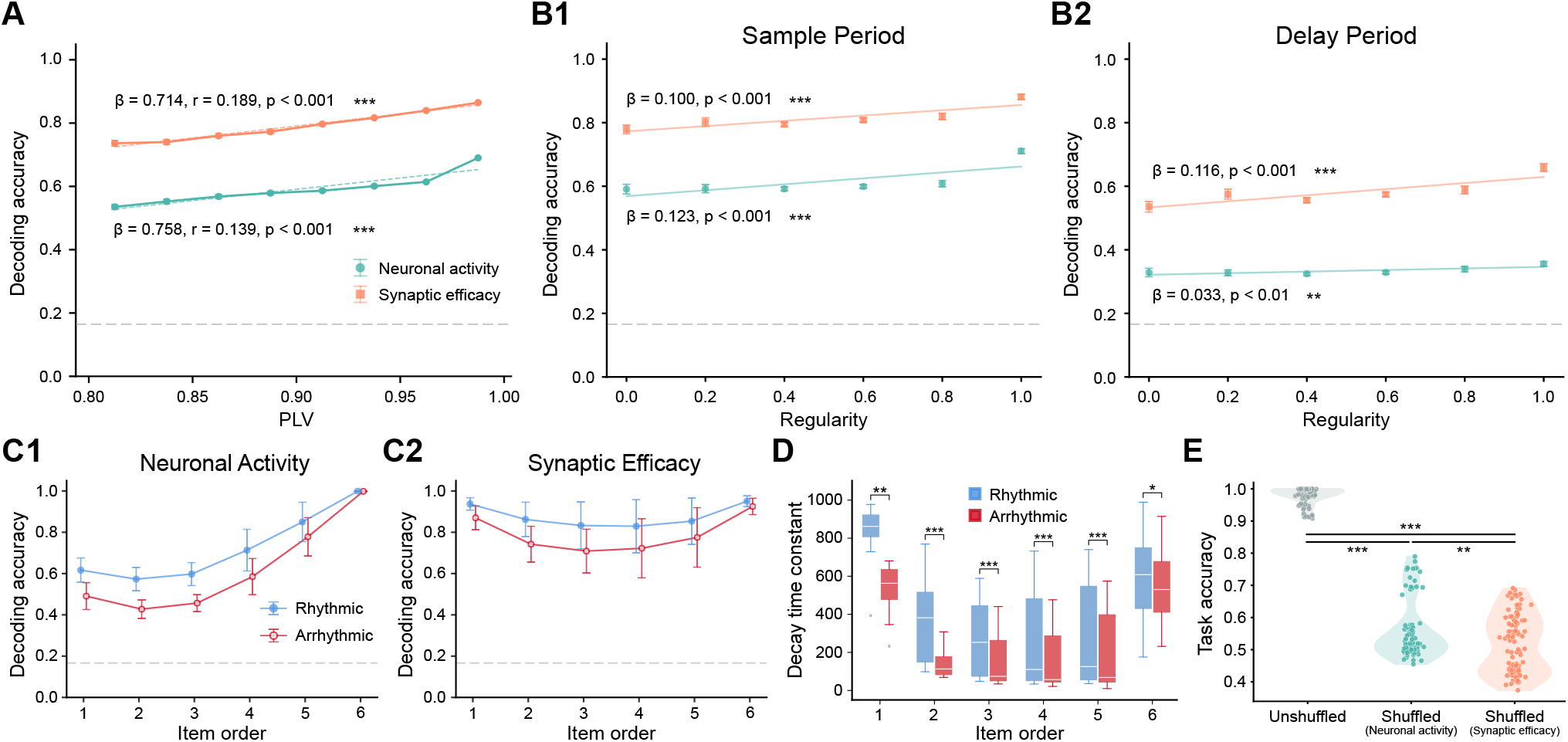
Rhythmic temporal structure enhances information decodability and prolongs decoding-derived memory persistence. (A) PLV-binned decoding accuracy analysis during the sample period. Item-level trials were grouped into fixed PLV intervals, and decoding accuracy was averaged within each bin. Higher PLV was associated with higher decoding accuracy for both neuronal activity and synaptic efficacy, consistent with an association between stronger phase locking and improved item-level information readout (Pearson correlation, two-sided, *n* = 6000 trials, *p* < 0.001). (B1-B2) Linear regression analysis linking decoding accuracy to the input regularity index (*R*) during the sample (Left) and delay (Right) periods. The significant positive correlations indicate that temporal regularity was associated with higher-fidelity stimulus information during both encoding and maintenance (Pearson correlation, two-sided, *n* = 6000 trials, *p* < 0.001). (C1-C2) Mean decoding accuracy for each sequence item (averaged across the sample and delay periods). At both the neuronal (C1) and synaptic (C2) levels, rhythmic input showed higher decoding accuracy across all 6 positions. This U-shaped pattern is consistent with classic serial-position effects. (D) Decay time constants (*τ*) of the decoding trajectories for the six items. For each independent network, decoding accuracy was estimated from 1000 trials and fitted with an exponential decay function to obtain item-specific *τ* values. Rhythmic input were associated with larger decay constants, indicating slower information decay. For each sequence item, rhythmic and arrhythmic conditions were compared across networks using a two-sided Wilcoxon signed-rank test with Bonferroni correction for six item-wise comparisons (*n* = 15 independent networks). (E) Impact of perturbing different network components on task accuracy. Violin plots show task accuracy under original conditions, shuffled neuronal activity, and shuffled synaptic efficacy. Shuffling synaptic efficacy caused a larger drop in accuracy compared to shuffling neuronal activity. Significance was assessed using a two-sided paired t-test with Bonferroni correction (*n* = 15 independent networks; corrected *p* < 0.01 and corrected *p* < 0.001). Unless otherwise stated, data are shown as mean ± s.e.m. Statistical tests are two-sided where applicable. Significance levels are denoted as \**p* < 0.05, \*\**p* < 0.01, and \*\*\**p* < 0.001.

Building on this global parametric enhancement, we examined how this rhythmic advantage is distributed across different sequence items (***Figure 5***C1 and ***Figure 5***C2). At both the neuronal and synaptic levels, rhythmic input was associated with higher mean decoding accuracy across all six positions, displaying a characteristic U-shaped serial position curve (primacy and recency effects) consistent with classical behavioral findings (***Orlov et al., 2000***; ***Hurlstone et al., 2014***). Notably, for intermediate items that are most susceptible to interference from subsequent stimuli (e.g., Item 2 and Item 3), the improvement brought by rhythmic input is particularly evident. Furthermore, by applying exponential fitting to the underlying temporal decay trajectories of the decoding accuracy, we estimated a memory decay time constant *τ* (***Figure 5***D) for each stimulus position. This decoding-based *τ* provides a population-level measure of information persistence, conceptually paralleling temporal context cells in macaque mPPC whose event-triggered responses decay with heterogeneous time constants during temporal order memory (***Zuo et al., 2026***). The decoding-based persistence of all six items was longer under the rhythmic condition, with some poorly remembered items showing a two-to nearly three-fold larger median *τ* than the arrhythmic condition (Item 2, 382 ± 59 vs. 137 ± 19, mean ± s.e.m.). This suggests that temporal regularity may slow the decay of decodable stimulus information in the network.

Finally, although the decoding analyses indicated robust retention of stimulus information, we asked whether synaptic efficacy made a functional contribution to working memory maintenance. We introduced perturbation experiments during the delay period by applying a random shuffle to either the neuronal activity or the synaptic efficacy (***Figure 5***E) (see Methods: Perturbation analysis). Under this perturbation protocol, shuffling synaptic efficacy produced a larger decline in task performance than shuffling neuronal activity (**, *p* < 0.01; ***, *p* < 0.001; two-sided paired t-test with Bonferroni correction, *n* = 15 independent networks), suggesting that synaptic efficacy contributed more to delay-period maintenance in the present model (***Masse et al., 2019***).

Together, these analyses are consistent with a cross-level account in which external regularity is associated with more stable phase organization, stronger temporally structured representations, and more persistent decodable memory traces.

## Discussion

This study examined how temporal regularity in external input facilitates sequential working memory in a biologically constrained recurrent neural network with excitatory-inhibitory structure and short-term synaptic plasticity. Rhythmic input improved sequence-memory performance and was accompanied by a more separable low-dimensional population geometry during encoding. Spectral and phase-based analyses further showed that the effect of temporal regularity was expressed around the dominant input frequency, with stronger stimulus-matched responses, reduced deviation between stimulus-arrival phase and response-favorable phase bins, and stronger locking between internal oscillatory phase and external temporal phase. Finally, time-resolved decoding and perturbation analyses indicated that rhythmic input improved the persistence of stimulus information in both neuronal activity and synaptic efficacy, with synaptic efficacy making a particularly important contribution during the delay period. Together, these results suggest that temporal regularity facilitates sequential working memory by organizing encoding-period recurrent dynamics and by supporting more stable synaptic-state representations during maintenance.

A central implication of these findings is that the rhythmic advantage cannot be explained solely by a nonspecific increase in response magnitude. Although temporal regularity increased activity in the band centered on the dominant input frequency, the phase analyses provided a more selective account of how rhythmic structure shaped network dynamics. Specifically, stimulus arrivals became progressively closer to phase bins associated with stronger responses, and this relationship was weak at the first item but strengthened as the sequence unfolded. This temporal profile suggests that the network did not merely respond independently to each stimulus onset. Instead, regular inter-onset intervals allowed neuronal activity to form a more stable relationship between external event timing and internal phase state. This interpretation is consistent with neurophysiological work showing that rhythmic stimulation can align low-frequency neural activity with expected event times and modulate sensory processing (***Lakatos et al., 2008***; ***Schroeder and Lakatos, 2009***; ***Nobre and Van Ede, 2018***). However, the present results should be interpreted as evidence for phase organization in a computational model rather than as direct evidence of biological entrainment. Phase locking can also reflect temporal predictability or repeated temporal preparation without necessarily demonstrating self-sustained entrainment (***Breska and Deouell, 2017***). For this reason, the current model is best understood as identifying a dynamical mechanism by which temporal regularity constrains recurrent phase states during sequence encoding.

The population-level findings further suggest that temporal regularity improves sequential memory by reducing variability in the timing of recurrent state evolution. Sequential working memory requires the network to encode multiple items while preserving their ordinal context, and therefore depends not only on representing item direction but also on separating successive item states. Under rhythmic input, task-related trajectories in temporal, ordinal, and stimulus-related subspaces were more clearly separated, indicating that regular timing helped organize population activity into more reproducible state trajectories. This interpretation is compatible with the broader view that neural population activity often evolves on low-dimensional manifolds whose geometry constrains downstream readout and task performance (***Cunningham and Yu, 2014***; ***Chung and Abbott, 2021***). In the present model, the enhanced geometric separation under rhythmic input may arise because stimulus-evoked responses are embedded within a more regular phase structure. By stabilizing when each item is encoded relative to the internal oscillatory cycle, temporal regularity may reduce overlap among adjacent item representations and thereby improve sequence-level readout.

Importantly, the representational consequences of oscillatory organization were not uniform across all task variables. Trial-level analyses showed that oscillatory power and phase-locking value were most consistently associated with selectivity for temporal-order variables, including stimulus order and interval order. In contrast, stimulus-direction selectivity did not show the same positive relationship with phase locking. This dissociation argues against a simple global-gain account in which rhythmic input uniformly sharpens all representations. Instead, rhythmic temporal structure preferentially supported aspects of the task that were intrinsically temporal. This result is theoretically important because serial-order information is a central challenge across verbal, visual, and spatial short-term memory domains (***Hurlstone et al., 2014***). It also provides a computational bridge to behavioral and electrophysiological findings showing that temporal structure at encoding can influence later recognition and that temporal expectations can dynamically prioritize working-memory representations for upcoming behavior (***Jones and Ward, 2019***; ***van Ede et al., 2017***; ***Jin et al., 2020***). The present model extends these findings by specifying how temporal regularity may shape recurrent population dynamics during sequence encoding.

The decoding and perturbation results link encoding-period organization to delay-period maintenance. Working memory has traditionally been associated with persistent neural activity. Still, a growing body of evidence indicates that information can also be retained in activity-silent or latent states that are not continuously expressed as elevated firing (***Stokes, 2015***; ***Rose et al., 2016***; ***Masse et al., 2019***; ***Barbosa et al., 2020***). Short-term synaptic plasticity provides one candidate mechanism for such latent maintenance, because transient changes in synaptic efficacy can preserve information over short timescales without requiring continuous activity (***Mongillo et al., 2008***; ***Masse et al., 2019***). In the present model, rhythmic input increased the decodability and persistence of stimulus information in both neuronal activity and synaptic efficacy. Moreover, disrupting synaptic efficacy during the delay impaired task performance more strongly than disrupting neuronal activity. These findings suggest that the benefit of rhythmic encoding extends beyond the sample period. Instead, more temporally organized encoding dynamics may leave more stable traces in synaptic variables, which subsequently contribute to delay-period maintenance.

Several limitations should be considered when interpreting the model. First, the recurrent units are simplified firing-rate units and should not be equated with biophysically detailed cortical neurons. Although the model incorporates excitatory–inhibitory structure and short-term synaptic plasticity, it does not capture the diversity of cortical cell types, laminar organization, neuromodulatory influences, or conductance-level synaptic mechanisms. Second, the oscillatory signatures reported here were extracted from model population activity; they should therefore be treated as dynamical regularities in the trained network rather than as direct equivalents of local-field-potential or EEG oscillations. Third, the model did not incorporate hierarchical cross-frequency interactions, such as theta–gamma coupling, which has been implicated in multi-item working memory (***Axmacher et al., 2010***). Future work should test whether incorporating cross-frequency mechanisms changes the relationship between temporal regularity, ordinal coding, and synaptic maintenance.

The model generates several empirically testable predictions. First, in biological sequential working-memory tasks, increasing the regularity of inter-onset intervals should strengthen low-frequency power and phase consistency near the dominant input frequency in task-relevant sensory and association areas. Second, the strength of phase organization near the end of the encoding period should predict the subsequent persistence of item information during the delay. Third, if synaptic-state variables contribute to the rhythmic advantage, perturbations that disrupt short-term facilitation or depression should reduce the behavioral benefit of rhythmic input, particularly when successful performance depends on delay-period maintenance. These predictions could be examined with high-temporal-resolution electrophysiological recordings, laminar recordings, or causal perturbation approaches in tasks that manipulate temporal regularity while holding item direction and memory load constant.

In summary, the present study provides a computational account of how rhythmic temporal structure facilitates sequential working memory. Temporal regularity organized recurrent population dynamics during encoding, aligned internal phase states with external event timing, preferentially supported temporal-order representations, and enhanced the persistence of stimulus information in synaptic efficacy during maintenance. These findings suggest that rhythmic input improves sequential working memory not simply by increasing neural responsiveness, but by imposing a more reliable temporal organization on the recurrent dynamics from which stable memory representations can emerge.

## Methods and Materials

### Network models

All simulations and training procedures were implemented using the PyTorch framework, with specific architecture and training parameters detailed in ***Table 1***. Task stimuli consisted of coherent motion patterns oriented along one of six discrete directions. The network architecture comprised an input layer of 31 units (30 direction-tuned neurons and 1 rule-tuned neuron) projecting to a recurrent layer of 100 neurons (***Barak, 2017***; ***Masse et al., 2019***), which in turn connected to three output units. Neurons within the recurrent layer were fully connected.

**Table 1.** Hyperparameters used for network architecture and training.

| Hyperparameter | Symbol | Value |
| --- | --- | --- |
| Learning rate | n/a | 0.02 |
| Neuron time constant | $\tau$ | 100 ms |
| Time step (training and testing) | $\Delta t$ | 10 ms |
| s.d. of input noise | $\sigma_{in}$ | 0.05 |
| s.d. of recurrent noise | $\sigma_{rec}$ | 0.25 |
| L2 penalty term on firing rates | $\lambda_{spike}$ | 0.01 |
| L2 penalty term on weights | $\lambda_{weight}$ | 0.001 |
| STSP neurotransmitter time constant | $\tau_x$ | 200 ms for facilitating<br>1500 ms for depressing |
| STSP utilization time constant | $\tau_u$ | 1500 ms for facilitating<br>200 ms for depressing |
| STSP neurotransmitter increment | $U$ | 0.15 for facilitating<br>0.45 for depressing |
| Number of neurons (input, recurrent, output) | $N_{in}, N_{rec}, N_{out}$ | 31, 100, 3 |
| Batch size | n/a | 1024 |
| Number of batches used to train network | n/a | 4000 |

The dynamics of the recurrent neural network were modeled using continuous-time firing rates. The activity *r*(*t*) of the recurrent units evolves according to the following differential equation:

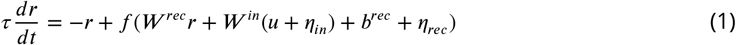

where *τ* denotes the neuronal time constant, *r* represents the firing rate vector of the recurrent layer, and *u* represents the input layer activity. The function *f* (⋅) serves as the non-linear activation function. *W* ^*rec*^ and *W* ^*in*^ denote the recurrent and input weight matrices, respectively, and *b*^*rec*^ is the recurrent bias vector.

To simulate trial-to-trial variability and neural noise, we injected independent Gaussian noise terms *η*_*in*_ and *η*_*rec*_ into the input and recurrent currents, respectively. These noise processes are defined as:

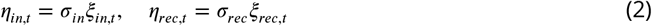

where *ξ* ∼ 𝒩 (0, 1) represents standard Gaussian white noise. To strictly enforce the biological constraint of non-negative firing rates, we employed the Rectified Linear Unit (ReLU) as the activation function:

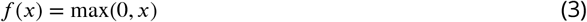

The neuronal activity is projected to the output layer via a linear transformation. To ensure numerical stability and compatibility with the Cross-Entropy loss function used during training, the model outputs *z* directly, without a softmax activation at the output layer:

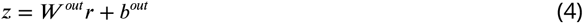

For numerical simulation, the continuous dynamics were discretized using a first-order Euler approximation with a time step of Δ*t*. The update rule is given by:

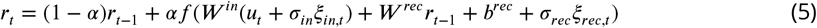

Because *f* (⋅) was implemented as a ReLU function and 0 < *α* ≤ 1, the recurrent firing rates remained non-negative, provided that the initial activity was non-negative.

To adhere to Dale’s law, the recurrent population was partitioned into 80 excitatory and 20 inhibitory units. We decomposed the recurrent weight matrix *W* ^*rec*^ into the product of a strictly non-negative matrix *W* ^*rec*+^ and a fixed diagonal matrix *D*:

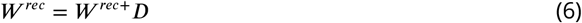

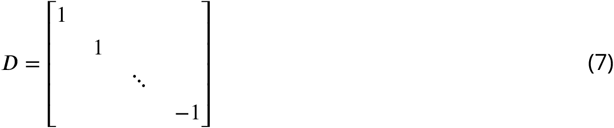

This formulation ensures that the first 80 neurons exert exclusively excitatory effects, while the remaining 20 exert inhibitory effects. Furthermore, self-connections were eliminated by enforcing all diagonal elements of *W* ^*rec*^ to be zero.

Unlike standard normal initialization, synaptic weights *W* ^*init*^ were sampled from a Gamma distribution (shape parameter *k* = 0.1, scale parameter *θ* = 1.0). This choice is motivated by prior theoretical work suggesting that Gamma-distributed connectivity facilitates faster convergence compared to uniform or Gaussian initialization in recurrent networks (***Masse et al., 2019***).

The input layer consists of 30 direction-selective neurons and one rule-encoding neuron. The 30 direction-selective units map discrete stimulus directions into continuous population activity patterns using von Mises tuning curves. The activity of input neuron *i* at time *t*, denoted as 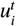, is defined as:

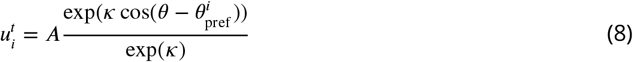

where *θ* represents the stimulus direction chosen from *K* = 6 possible directions 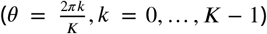. Each neuron *i* has a preferred direction 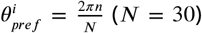. The tuning parameters were set to *κ* = 2 (concentration) and *A* = 4 (amplitude) during stimulus presentation (sample and test epochs). During non-stimulus epochs (fixation, delay, and response), the amplitude *A* was set to 0. Additionally, a binary rule-encoding unit was included to indicate task epochs. This unit was set to active (1) during fixation and delay requirements, and inactive (0) during the response epoch (match/non-match decision). The specific scaling and count of this rule unit are arbitrary and do not significantly influence the network’s training dynamics.

### Network training

We trained the network parameters using supervised learning via the Backpropagation Through Time (BPTT) algorithm (***Werbos, 2002***). The total objective function, *L*_*total*_, comprised three components: a task performance loss, a metabolic spike penalty, and a synaptic weight penalty. The training objective was to minimize: (1) the Cross-Entropy between the network output and the target sequence, which forces the network to produce correct decisions at the correct timing; (2) the Firing Rate Regularization of hidden units, which enforces a sparse coding strategy to mimic biological metabolic constraints and prevent unrealistic high-frequency discharge; and (3) the Connection Weight Regularization, which maintains structural sparsity and prevents dynamic instability caused by excessive synaptic weights.

The target output was represented as a time-varying one-hot vector of dimension 3. The first unit encoded the ‘fixation/wait’ state (active during fixation, sample, delay, and test epochs). During the response epoch, the second unit encoded a ‘match’ decision, while the third unit encoded a ‘non-match’ decision. The specific loss components are defined as follows:

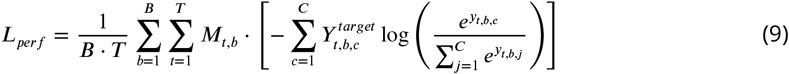

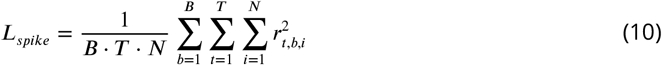

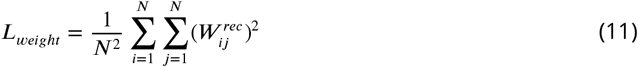

where *B* denotes the batch size, *T* is the total number of time steps in a trial, and *N* represents the number of recurrent neurons. *M*_*t,b*_ is a temporal loss mask, and *r*_*t,b,i*_ represents the firing rate of neuron *i*. The total loss is a weighted sum:

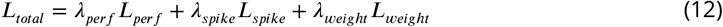

The weighting coefficients were set to *λ*_*perf*_ = 1.0, *λ*_*spike*_ = 0.01, and *λ*_*weight*_ = 0.001.

To prioritize decision accuracy, we applied a dynamic weighting scheme to the loss mask *M*_*t,b*_. Specifically, the loss weight for the response period (*λ*_*res*_) was set to 2.0, while non-response periods were weighted at 1.0 (*λ*_*non*_*res*_). Additionally, we implemented a 20 ms “grace period” at the onset of the response period, during which the loss mask was set to 0, allowing the network sufficient time to transition into the decision state.

Network parameters were optimized using the Adam optimizer (***Kingma and Ba, 2014***). To ensure smooth convergence, we employed a StepLR learning rate scheduler, decaying the learning rate by 50% every 100 epochs. To mitigate the exploding gradient problem common in recurrent neural networks, we applied gradient clipping, restricting the global norm of gradients to 1.0 (***Pascanu et al., 2013***).

Model performance was evaluated based on sequence-level accuracy. A trial was considered correct only if the argmax of the network’s output probability distribution matched the target class at every time step throughout the task (excluding the 20 ms grace period). This strict “full-sequence matching” criterion ensures the reliability of the performance metric by penalizing transient instabilities or random guessing.

### Short-term synaptic plasticity

To incorporate biologically realistic memory mechanisms, the synaptic efficacy between recurrently connected neurons was dynamically modulated via short-term synaptic plasticity (STSP). Following the phenomenological framework established by ***Mongillo et al. (2008***), we modeled STSP through the interaction of two state variables: *x*(*t*), representing the fraction of available neurotransmitter resources, and *u*(*t*), representing the utilization of those resources. Presynaptic activity *r*(*t*) drives the influx of calcium into the presynaptic terminal, which increases the utilization *u*(*t*) and facilitates synaptic transmission. Simultaneously, each presynaptic spike depletes the available neurotransmitter pool *x*(*t*), leading to synaptic depression. The temporal evolution of these variables is governed by the following differential equations:

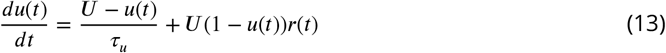

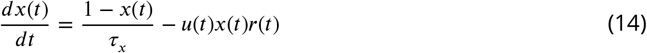

where *U* is the baseline utilization level, *τ*_*u*_ is the decay time constant for facilitation, and *τ*_*x*_ is the recovery time constant for depression. The net synaptic input *J* (*t*) received by a postsynaptic neuron is the product of the fixed synaptic weight *W* and the instantaneous values of *u*(*t*) and *x*(*t*):

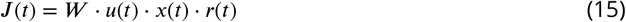

In our network, 50% of the recurrent neurons were assigned facilitating synapses (STF), characterized by a utilization time constant *τ*_*u*_ significantly longer than the recovery time constant *τ*_*x*_. The remaining 50% of neurons projected depressing synapses (STD), where *τ*_*x*_ dominated over *τ*_*u*_. For computational efficiency, these STSP variables were updated at the neuronal level, such that all outgoing synapses from the same presynaptic unit shared identical dynamics. STSP was applied exclusively to recurrent connections, while input and output weights remained static.

### Task paradigm

We employed a sequential delayed match-to-sample (DMS) task to investigate the impact of temporal rhythmicity on working memory performance. Each trial commenced with a 500 ms fixation period, followed by a sample period where a sequence of six oriented stimuli was presented. Stimulus orientations were drawn from a set of six angles (0° to 300° in 60° increments), with the constraint that adjacent stimuli in the sequence always differed in orientation.

To isolate the effect of temporal structure, we designed two experimental conditions for the sample period: a rhythmic condition and an arrhythmic condition. These temporal parameters were strictly constrained by classical behavioral and psychophysical findings. Empirical evidence indicates that the optimal psychological integration window for rhythmic sequences typically spans 1 to 5 seconds, with human beat perception functioning most reliably across inter-onset intervals (IOIs) ranging from 100 to 1500 ms (***Krumhansl, 2000***). Furthermore, the preferred processing rate for sequential temporal events is approximately 0.8 to 6 items per second (***Halpern and Müllensiefen, 2008***; ***Warren et al., 1991***). Accordingly, in the rhythmic condition, stimuli were presented with a constant inter-onset interval (IOI) of 250/300/350/400/450/500 ms and a constant duration of 100/120/140 ms. In the arrhythmic condition, while the total duration of the sample sequence was matched to the rhythmic condition (falling within the 1-5s integration capacity (***Levitin et al., 2018***)), the IOIs were jittered randomly between 40% and 160% of the mean IOI (***Tian et al., 2025***). Following a 1000 ms delay period, a test sequence was presented. In match trials, the test sequence was identical to the sample; in non-match trials, the orientations of either the odd-indexed (1st, 3rd, 5th) or even-indexed (2nd, 4th, 6th) stimuli were rotated by 60°. Crucially, to ensure that any performance differences arose solely from the encoding period, the temporal structure of the test sequence was “semi-rhythmic” (IOIs jittered between 70% and 130%) and identical across all trials for a given model instance.

Finally, a response cue was presented for 100 ms, prompting the model to report whether the sample and test sequences matched. The dataset was balanced across conditions, comprising 18,750 match and 37,500 non-match unique orientation sequences, each instantiated in both rhythmic and arrhythmic temporal formats. During training, batches were stratified to contain equal numbers of rhythmic-match, rhythmic-non-match, arrhythmic-match, and arrhythmic-non-match trials. To verify the robustness of our findings, we systematically varied the stimulus duration (100–140 ms) and mean IOI (250–500 ms) across independent experimental runs.

To quantitatively evaluate the degree of temporal rhythmicity for individual trials across all conditions, we computed a trial-wise regularity index *R*. First, the coefficient of variation of the interonset intervals *CV*_*IOI*_ was calculated as the ratio of a single trial’s standard deviation to the mean 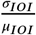. We then defined the regularity index *R* by applying Min-Max normalization to the quantity 1 − *CV*_*IOI*_ across the entire dataset. This transformation yields a strictly bounded metric (*R* ∈ [0, 1]), where *R* = 1 denotes perfect periodicity and lower values indicate increasing temporal jitter.

### Demixed principal component analysis

To capture the shared dynamical features of neural trajectories during the memory encoding process across varying trials and conditions, we performed temporal alignment on the hidden layer activity. Given the trial-to-trial variability of Inter-Onset Intervals (IOIs) across different models, neural activity corresponding to each stimulus position (Positions 1–6) was extracted and time-warped to a fixed duration of 400 ms using linear interpolation.

Consequently, the activity for each trial was segmented into six matrices, each denoted as 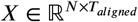, where *N* represents the number of recurrent neurons and *T*_*aligned*_ denotes the number of aligned time steps (spanning from one stimulus onset to the next). Prior to dimensionality reduction, all neural data were centered by subtracting the global mean activity.

To decompose the population activity into independent task-related latent components (temporal component, ordinal component, and directional component), we employed a marginalization procedure. For each neuron, we computed the marginal mean trajectories by averaging activity across specific conditions:

1. Temporal component: Averaged across rhythmic and arrhythmic conditions.
2. Ordinal component: Averaged across stimulus ordinal positions (1–6).
3. Directional component: Averaged across stimulus feature IDs (0–5).

These marginal means were then tiled to match the dimensions of the original data, yielding marginalized data matrices *X*_*ϕ*_ (where *ϕ* ∈ {*time, pos, stim*}). Unlike standard PCA, demixed PCA (dPCA) seeks a low-rank subspace that maximizes the variance explained by these specific marginalizations, decomposing the total variance into (***Kobak et al., 2016***):

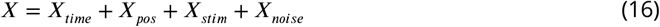

This formulation isolates task-specific components from residual noise. We solved for the decoding matrix *D* using reduced-rank regression (RRR). To ensure numerical stability and prevent overfitting during the inversion of the covariance matrix, a ridge regularization term (*λ* = 10^−6^) was introduced:

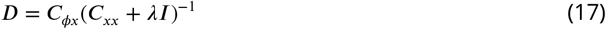

where *C*_*xx*_ is the covariance matrix of the original data, and *C*_*ϕx*_ is the cross-covariance matrix between the marginalized components and the original data. Principal Component Analysis (PCA) was then performed on the predicted activity 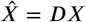 to extract the top five distinct components. Each extracted dPCA component *w* was automatically assigned to the marginalization *ϕ* that explained the largest fraction of its variance:

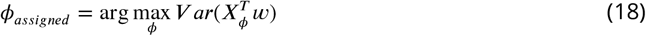

To assess the robustness of the trajectories, this analysis was replicated across 10 independent models. We report the mean projected trajectories, with shaded regions indicating the standard error of the mean (s.e.m.). To quantify the representational efficacy of the extracted components, we defined two variance metrics:

1. Global Variance Ratio (*R*_*g*_):

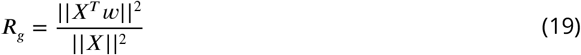 This metric measures the component’s contribution to the total neural fluctuation (including noise and task-irrelevant dynamics). Due to the high dimensionality and noise inherent in neural networks, this value is typically small.
2. Marginal Variance Ratio (*R*_*m*_):

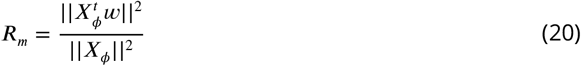

This metric quantifies the component’s dominance within the purified, task-relevant subspace. A high value indicates that the component serves as a primary feature for that specific task dimension, signifying effective signal isolation.

### Trial-wise temporal regularity and stimulation frequency

Trial-wise temporal regularity was quantified from the sequence of sample stimulus onset times. Let *s*_*r,j*_ denote the onset time of the *j*-th stimulus in trial *r*. The inter-onset intervals were defined as

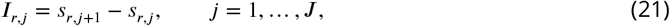

where *J* = 5 for a sequence of six sample stimuli. The mean inter-onset interval of trial *r* was

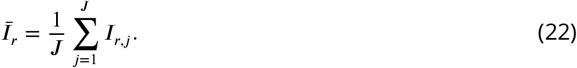

Raw temporal regularity was defined as

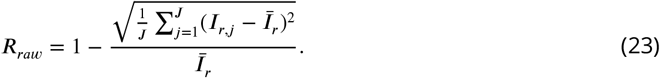

Then, we defined the regularity index *R* as Min-Max normalized *R*_*raw*_ across the entire dataset. Thus, *R* = 1 indicates perfectly periodic stimulation, whereas smaller values indicate greater temporal variability. For binned analyses, non-periodic trials were divided into quantile-matched regularity bins, whereas perfectly periodic trials formed a separate bin.

The characteristic stimulation frequency was defined as

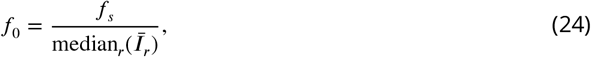

where *f*_*s*_ = 100 Hz is the sampling rate, and the median was taken across trials.

### Population signal construction for oscillatory analysis

For each trial, population activity during the sample epoch was represented as a neuronal state vector **x**_**r**_(*t*) ∈ ℝ^*N*^, where *N* denotes the number of units. To obtain a robust population-level signal for subsequent spectral and phase analyses, neural activity was standardized across units to remove differences in activity scale and then projected onto principal components. This step reduces unit-specific variability while preserving the dominant temporal structure shared across the population.

Principal components were retained until cumulative explained variance exceeded 90%. A scalar population signal was then constructed as the explained-variance-weighted projection of the retained component scores,

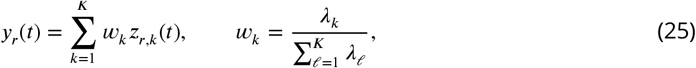

where *z*_*r,k*_(*t*) is the score of the *k*-th principal component in trial *r, λ*_*k*_ is the explained variance of that component, and *K* is the number of retained components.

### Spectral analysis of population activity

The time–frequency structure of the population signal *y*_*r*_(*t*) was characterized using complex Morlet wavelets,

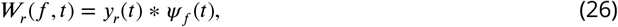

with

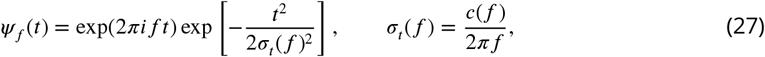

where *σ*_*t*_(*f*) is the temporal standard deviation of the wavelet and *c*(*f*) denotes the number of cycles. Each wavelet was normalized to unit energy before convolution. Power was computed as

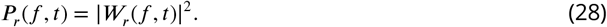

Sample-period spectral power was obtained by averaging power across the sample epoch,

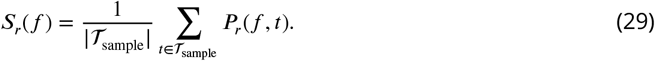

To isolate narrowband oscillatory responses from broadband spectral structure, spectra were log-transformed and detrended along the relative-frequency axis *m* = *f* /*f*_0_,

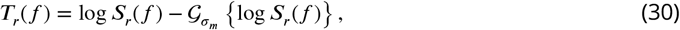

where 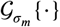 denotes Gaussian smoothing over *m*, and *σ*_*m*_ is the smoothing width in units of relative frequency.

Stimulus-responsive frequencies were tested using a one-sample sign-flip cluster permutation test against zero. At each frequency, the observed statistic was

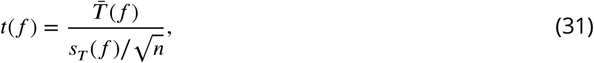

where 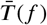 and *s*_*T*_ (*f*) denote the across-trial mean and standard deviation of *T*_*r*_(*f*), and *n* is the number of trials. Adjacent frequencies exceeding a one-sided cluster-forming threshold were grouped into clusters, with cluster mass defined as

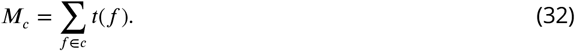

The null distribution was obtained by randomly multiplying each trial spectrum by +1 or −1 and retaining the maximum cluster mass across frequencies for each permutation. Cluster-level significance was computed as

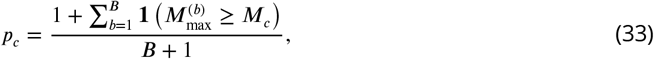

where *B* = 1000 is the number of permutations and **1**(⋅) is the indicator function. Effect size was reported as Cohen’s *d*_*z*_,

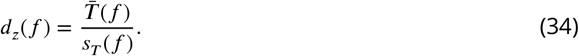

Regularity-dependent spectral modulation was tested using a frequency-wise one-way analysis of variance across regularity bins. For each frequency,

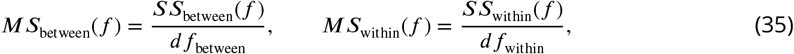

and

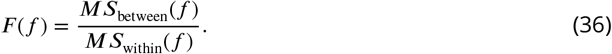

Cluster-level inference was performed by permuting regularity-bin labels across trials and recomputing the maximum suprathreshold cluster mass. Effect size was quantified using omega squared,

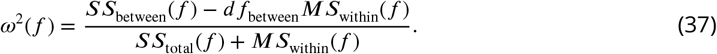

The direction of regularity modulation was characterized using Spearman’s rank correlation,

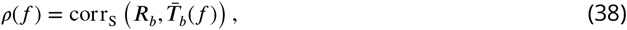

where *R*_*b*_ is the median regularity of bin *b*, 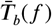 is the mean detrended response within that bin, and corr_S_ denotes Spearman’s rank correlation.

Candidate frequency components were evaluated around stimulation-related frequencies 0.5*f*_0_, 1*f*_0_, 2*f*_0_, and 3*f*_0_. A component was retained as a primary frequency component if it showed both significant stimulus responsiveness and significant regularity modulation, contained a positive local spectral peak, and exhibited non-negative monotonic dependence on regularity. Components showing significant stimulus responsiveness without significant regularity modulation were retained as secondary response components. Final analysis bands were defined as

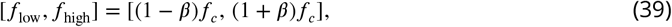

where *f*_*c*_ is the selected center frequency and *β* = 0.20.

### Oscillatory phase organization during sample encoding

Phase analyses were performed on a low-dimensional population signal extracted from recurrent activity. For each analysis condition, trial activity was first projected onto principal components after robust preprocessing, including percentile clipping, z-scoring with a minimum standard-deviation floor, and z-score clipping. The number of retained components was selected to explain at least 90% of the variance. A one-dimensional population signal was then constructed as the explained-variance-weighted sum of the retained principal-component scores:

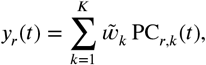

where *K* is the number of retained components and 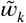 denotes the normalized explained-variance weight of component *k*.

For phase analyses, this population signal was transformed into a complex analytic-like signal using a causal complex filter within the retained frequency band. Let [*f*_low_, *f*_high_] denote the selected band around the stimulation-related frequency component, and let *f*_*c*_ = (*f*_low_ + *f*_high_)/2. The causal complex kernel was defined over non-negative lags as

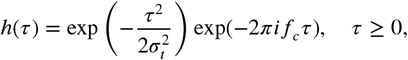

with the support truncated to a maximum of 0.45 s. The resulting complex signal was

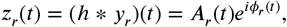

where *A*_*r*_(*t*) = *z*_*r*_(*t*) is the instantaneous band amplitude and *ϕ*_*r*_(*t*) = arg[*z*_*r*_(*t*)] is the instantaneous phase. The sign of the unwrapped phase was adjusted so that phase progression was positive in perfectly regular trials.

For each trial *r* and stimulus position *j*, the stimulus-arrival phase was extracted at the onset time *s*_*r,j*_ :

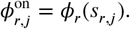

To reduce unreliable phase estimates, trials with low onset-band amplitude were excluded separately for each stimulus position. Specifically, a trial was retained for position *j* only if *A*_*r*_(*s*_*r,j*_) was above the 25th percentile of valid onset amplitudes for that position.

A response-favorable phase was estimated separately for each stimulus position. We first computed a local post-onset population-response metric *M*_*r,j*_ within the stimulus window:

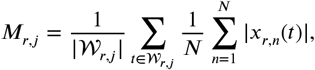

where 𝒲_*r,j*_ = [*s*_*r,j*_, *s*_*r,j*_ + *L*_stim_), *x*_*r,n*_ (*t*) is the activity of recurrent unit *n*, and *L*_stim_ is the stimulus duration. Stimulus-arrival phases were then binned into 10-degree bins. For each phase bin, we computed the mean local response *M*_*r,j*_, smoothed the phase-response profile circularly, and multiplied it by a density-support term derived from the number of samples in that bin. The resulting density-regularized response score was used to define the preferred encoding phase:

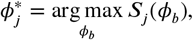

where *S*_*j*_(*ϕ*_*b*_) is the smoothed density-regularized response score for phase bin *ϕ*_*b*_. Thus, 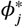 denotes the stimulus-arrival phase associated with the strongest local population response for stimulus position *j*, rather than the circular mean phase of the stimulus window.

The deviation between the stimulus-arrival phase and the preferred encoding phase was quantified as the absolute circular distance:

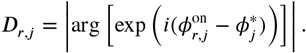

Smaller *D*_*r,j*_ indicates that the stimulus arrived closer to the response-favorable phase. For sequence-level summaries, deviations or alignment scores were averaged across stimulus positions, with the first onset excluded from the main sequence-level summary because it occurred before temporal context had accumulated. To quantify how stimulus presentation modulated phase progression, we estimated phase advancement within each stimulus window. For each trial *r* and stimulus position *j*, the signed unwrapped phase trajectory *ψ*_*r*_(*t*) was fitted within the stimulus window 𝒲 _*r,j*_ = [*s*_*r,j*_, *s*_*r,j*_ + *L*_stim_) using an amplitude-weighted robust linear model:

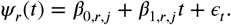

The fitting weights were derived from instantaneous band amplitude:

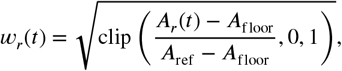

where *A*_floor_ and *A*_ref_ were defined as the 20th and 90th percentiles, respectively, of sample-period band amplitudes. The slope was estimated using an iterative robust weighted regression with Huber reweighting. Fits were retained only when the number of valid weighted samples exceeded the minimum valid-point criterion and the fitted trajectory exceeded a minimum *R*^2^ threshold.

The observed phase advance during the stimulus window was defined as

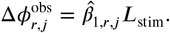

The nominal phase advance expected from the retained *f*_0_ component was

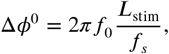

where *f*_*s*_ is the sampling rate. Phase kick was then defined as the circular difference between the observed and nominal phase advance:

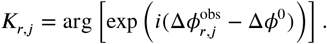

Positive values indicate faster-than-*f*_0_ phase advancement during stimulus presentation, whereas negative values indicate slower-than-*f*_0_ advancement.

To examine whether stimulus-induced phase advancement depended on the phase state at stimulus arrival, we related *K*_*r,j*_ to the signed stimulus-arrival phase offset from the preferred phase:

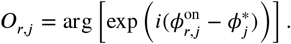

For visualization, trials were grouped into four regularity groups: low *R*, mid-low *R*, mid-high *R*, and perfectly regular trials (*R* = 1). The signed phase offsets were binned, and phase kick was averaged within each bin when the bin contained a sufficient number of trials.

### Inter-stimulus phase-progression frequency

To examine how internal oscillatory phase continued to advance between successive sample stimuli, we quantified phase progression during the inter-stimulus interval (ISI) following each stimulus. For trial *r* and transition *j*, the ISI window was defined as the interval from the offset of the current stimulus to the onset of the next stimulus:

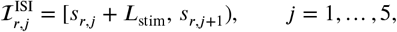

where *s*_*r,j*_ denotes the onset time of stimulus *j, L*_stim_ is the stimulus duration, and *s*_*r,j*+1_ denotes the onset time of the following stimulus.

Let *ϕ*_*r*_(*t*) denote the instantaneous phase of the retained *f*_0_-band population signal. To estimate the rate of phase advancement within each ISI window, the phase time series was first unwrapped to obtain a continuous phase trajectory *ψ*_*r*_(*t*). We then fitted a linear phase-progression model within each ISI window:

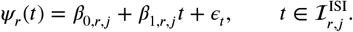

The fitted slope 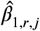 provides a window-level estimate of the average phase-advance rate. This slope was converted into an ISI phase-progression frequency:

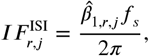

where *f*_*s*_ is the sampling rate. Thus, *IF* ^ISI^ summarizes how rapidly the internal oscillatory phase advanced between the offset of stimulus *j* and the onset of stimulus *j* + 1.

To quantify how far this ISI phase progression deviated from the dominant input timescale, we computed the absolute frequency deviation from *f*_0_:

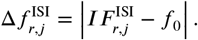

Here, smaller values indicate that the internal phase advanced at a rate closer to the dominant input frequency, whereas larger values indicate a greater deviation of ISI phase progression from the input-defined temporal scale.

For visualization, 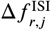 was averaged within trial-wise temporal-regularity bins for each transition. Continuous relationships between temporal regularity and ISI frequency deviation were assessed using Spearman’s rank correlation separately for each transition. A sequence-level summary was obtained by averaging 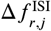 across later ISI windows after the first transition, and the same correlation analysis was applied to this averaged measure.

### Coupling between external temporal phase and neural oscillatory phase

To measure how strongly internal phase dynamics tracked the external temporal structure of the sequence, we constructed an external temporal phase for each inter-onset interval. For trial *r* and interval *j*, bounded by consecutive stimulus onsets *s*_*r,j*_ and *s*_*r,j*+1_, the external phase was defined as

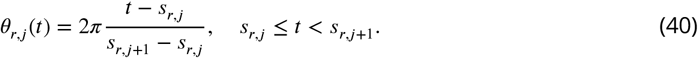

Thus, *θ*_*r,j*_ (*t*) advanced linearly from 0 to 2*π* within each inter-onset interval and reset at the next stimulus onset.

The phase difference between internal oscillatory phase and external temporal phase was computed as

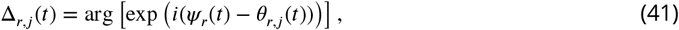

where *ψ*_*r*_(*t*) is the signed unwrapped internal phase used in the phase-slope analyses.

Phase locking was quantified using an amplitude-weighted phase-locking value:

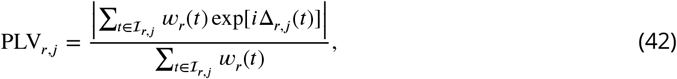

where ℐ_*r,j*_ = [*s*_*r,j*_, *s*_*r,j*+1_) and *w*_*r*_(*t*) is the same amplitude-derived weight used in the phase-slope analyses. PLV_*r,j*_ ∈ [0, 1], with larger values indicating a more consistent phase relationship between internal oscillations and the external temporal phase. The dependence of phase locking on temporal regularity was quantified for each interval using Spearman’s rank correlation,

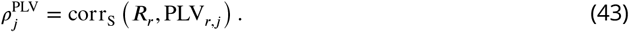

An overall phase-locking index was defined as the average PLV across inter-onset intervals,

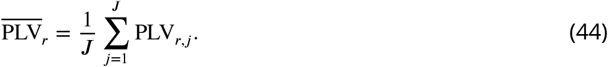

The relationship between 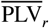 and trial-wise temporal regularity was assessed using the same permutation-based Spearman correlation procedure.

### Selectivity analysis of oscillatory organization

To examine how sample-period oscillatory organization was related to the encoding of different task variables, we combined unit-level selectivity estimates with unit-level oscillatory metrics across regularity bins. Trials were grouped according to the trial-wise temporal regularity index described above. For each independent model and each regularity bin, recurrent activity was used to estimate the selectivity of each hidden unit for three task variables: stimulus order, interval order, and stimulus direction. Before selectivity estimation, recurrent activity was z-scored across time for each unit.

For each unit, selectivity was quantified as the fraction of response variance explained by a task variable,

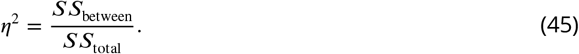

Here, *SS*_between_ denotes the variance explained by differences between task groups, and *SS*_total_ denotes the total response variance across all observations. Statistical significance of selectivity was assessed using a one-way ANOVA across task groups. Stimulus-order selectivity was computed from the mean activity within each stimulus-presentation window, grouped by the ordinal position of the sample stimulus (1, …, 6). Interval-order selectivity was computed from the mean activity within a fixed pre-onset segment of each inter-stimulus interval, grouped by interval index (1, …, 5). Stimulus-direction selectivity was computed from stimulus-window responses grouped by stimulus identity. To reduce confounding between direction identity and serial position, position-specific mean responses were subtracted before estimating direction selectivity.

For the same units and regularity bins, oscillatory organization was quantified within the retained stimulus-matched frequency band centered on *f*_0_. For unit *i* and regularity bin *b*, let *x*_*r,i*_(*t*) denote the activity of unit *i* in trial *r*, and let ℛ denote the set of trials assigned to bin *b*. The bin-averaged activity was first computed as

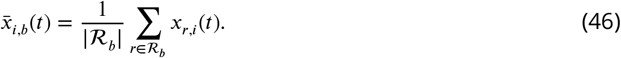

This bin-averaged activity was transformed into a complex band-limited signal using the same retained *f*_0_-band filtering procedure described above,

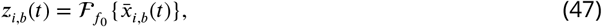

where 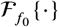 denotes the complex filter within the retained stimulus-matched frequency band. Oscillatory power was then defined as the mean-squared amplitude of this complex band-limited signal during the sample period,

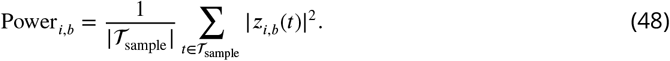

Unit-wise PLV was computed using the external temporal phase in Eq. 40 and the amplitude-weighted phase-locking definition in Eq. 42, replacing the population-level phase with the retained-band phase of each unit and averaging PLV within each regularity bin. Thus, both power and PLV entered the heatmap analysis as unit-by-regularity-bin measures.

To construct the joint heatmaps, units showing significant selectivity for a given task variable were first selected according to the ANOVA *p* value. These units were then assigned to equal-width *η*^2^ bins between 0 and 1. For each task variable and oscillatory metric, each heatmap cell corresponded to one regularity bin and one *η*^2^ bin. The cell value was computed as the mean power or PLV across all selected units falling into that bin combination within a model. This procedure yielded one matrix for each model, task variable, and oscillatory metric. The displayed heatmaps were obtained by aggregating the corresponding matrices across independent models using the median.

To quantify monotonic trends along the two heatmap dimensions, Spearman’s rank correlations were computed separately for each model across valid heatmap cells. The regularity-axis correlation measured whether the oscillatory metric varied monotonically with the regularity-bin coordinate,

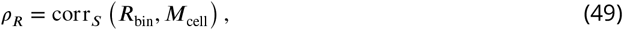

whereas the selectivity-axis correlation measured whether the oscillatory metric varied monotonically with the *η*^2^-bin coordinate,

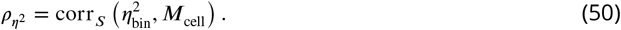

Here, *M*_cell_ denotes the heatmap value for power or PLV in a valid cell. Each model therefore contributed one *ρ*_*R*_ and one 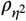 for each task variable and oscillatory metric. Group-level significance was assessed across independent models. Model-level Spearman correlations were first transformed using Fisher’s *z* transformation, and the resulting values were tested against zero using a two-sided sign-flip test.

### Population decoding

We quantified the network’s capacity to represent sequence information by measuring the accuracy of decoding stimulus identities using linear multi-class Support Vector Machines (SVMs). A linear classifier was chosen because the network’s readout layer inherently performs a linear classification on the recurrent population activity, a standard approach for assessing dynamic mental representations (***King and Dehaene, 2014***).

We implemented a time-resolved population decoding analysis using a sliding window approach. At each time step (10 ms stride), a linear classifier was trained on the activity of the 100 recurrent units to decode the direction of stimuli presented at specific ordinal positions (1–6) within the sequence. Decoding was performed independently for two distinct state variables:

1. Neural Activity: Defined as the instantaneous firing rates of the recurrent population.
2. Synaptic Efficacy: Defined as the product of available neurotransmitter resources *x*(*t*) and utilization fraction *u*(*t*) (i.e., *x*(*t*)⋅*u*(*t*)), representing the instantaneous synaptic strength as described above.

A Linear Support Vector Machine (Linear SVC) with a regularization parameter *C* = 1.0 served as the decoder. Decoding accuracy was assessed using 5-fold cross-validation on a dataset of 1000 trials per model.

To prevent data leakage and ensure rigorous evaluation, feature standardization (Z-score normalization) was computed solely on the training set within each fold and subsequently applied to the test set using the same parameters. This procedure normalized the scale of neuronal activity while preserving the independence of the test data. Performance was quantified as the proportion of correctly classified trials in the test set (80 trials per fold). The final reported accuracy represents the average across all five folds. Given the six possible stimulus categories, the theoretical chance level was 1/6 ≈ 16.7%.

To link phase locking with information readout, item-specific PLV values were paired with the decoding accuracy of the corresponding stimulus item. For the PLV-binned analysis, trials were grouped according to PLV, and decoding accuracy was averaged within each bin. For the regularity-dependent analysis, trial-level regularity *R* was paired with item-level decoding accuracy from the same trial. Correlations and linear regressions were computed from the unbinned observations, whereas plotted points show bin-averaged means. The reported slope denotes the unstandardized linear regression slope.

To visualize memory decay dynamics as a function of time, we derived six separate decoding curves corresponding to the stimuli at the six ordinal positions in the sample period. To aggregate results across 15 models trained with varying temporal structures (different IOIs and SDs), we employed a piecewise temporal resampling strategy to map all decoding trajectories onto a standardized timeline (Target IOI = 300*ms*, Target SD = 100*ms*). Specifically, decoding traces during the sample period were linearly warped (compressed or stretched) between consecutive stimulus onsets to align with the standardized epoch durations. This alignment ensured that task events were synchronized across models despite differences in their native temporal scales. The visualized trajectories represent the mean decoding accuracy averaged across the 15 models, with shaded regions indicating the standard error of the mean (s.e.m.).

To quantify the persistence of memory representation, we used the decay stage of the decoding curve (the sample period in the match dataset before the same stimulus was presented in the test period), and fitted it with an exponential decay function:

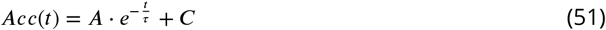

where *τ* represents the decay time constant, *A* is the amplitude, and *C* is the baseline chance level (1/6).

### Perturbation analysis

To examine whether task performance depended on the neuron-specific organization of internal network states during memory maintenance, we performed perturbation analyses on trained networks. During forward propagation, perturbations were applied at the start of the delay period. In the synaptic-state perturbation, the available neurotransmitter fraction *x* and utilization fraction *u* were independently shuffled across recurrent units for each trial. This manipulation preserved the population-level distribution of each synaptic variable at the perturbation time point, but disrupted its neuron-specific assignment and the local pairing between *x* and *u*. In the neuronal-state perturbation, hidden activity *h* was shuffled across recurrent units using the same procedure, preserving the distribution of activity values while disrupting their unit-specific assignment.

The external input sequence, recurrent weights, readout weights, and task labels were kept unchanged in all perturbation conditions. After perturbation, the network dynamics continued to evolve normally according to the same recurrent and STSP update equations. Thus, performance changes quantified the impact of disrupting structured synaptic or neuronal states carried by the trained recurrent network.

## Data availability

The simulation code, task-generation scripts, model-training scripts, and analysis code used in this study are publicly available at https://github.com/Bold2023/rhythmic_working_memory.

